# Distinct rich and diverse clubs regulate coarse and fine binocular disparity processing: Evidence from stereoscopic task-based fMRI

**DOI:** 10.1101/2023.10.25.564094

**Authors:** Kritika Lohia, Rijul Saurabh Soans, Rohit Saxena, Kabir Mahajan, Tapan K. Gandhi

## Abstract

While cortical regions involved in processing binocular disparities have been studied extensively, little is known on how the human visual system adapts to changing disparity magnitudes. Even though there is an established correlation of BOLD signal with disparity magnitudes, this correlation is not inherent and instead arises from specific causal interactions within an integrated network. Here, we investigate causal mechanisms of coarse and fine binocular disparity processing using fMRI with a clinically validated, custom anaglyph-based stimulus. Therefore, we use degree (D) and participation coefficient (PC) metrics representing rich and diverse properties of the brain network, respectively. Twenty-six healthy participants were asked to indicate hidden 3D shapes through anaglyph filters at four disparity magnitudes. Our findings reveal significant changes at different disparity magnitudes in terms of D and PC of Middle Temporal (MT), V2, V3 and Superior Parietal Lobule (SPL) across both hemispheres. Of these, MT exhibited overlapping rich and diverse club characteristics among other brain regions. Further, diverse clubs outperform rich clubs in decoding disparity magnitudes irrespective of the hemisphere, thereby reinforcing their integrative network properties. These findings imply that distinct rich and diverse clubs exist and provide functional evidence for the variability in human stereopsis.

## 1. Introduction

Stereopsis is a fundamental feature of the human visual system (HVS) that is essential for the reconstruction of the depth dimension of the world. The HVS is able to compute this due to the horizontal separation of the eyes that introduce tiny horizontal differences in the retinal images of objects. These differences – termed binocular disparities – are the first step towards the evaluation of stereopsis(1). Consequently, understanding the underlying neural mechanisms of binocular disparities has important implications – particularly, in the assessment and treatment of eye disorders such as strabismus and amblyopia(2–11), interaction with virtual reality(12,13), and inverse problems in computer vision(14–16).

Previous investigations on binocular disparities have revealed the presence of disparity-specific regions across dorsal and ventral visual streams including V3A, MT+/V5, V7, lateral occipital (LO) and intraparietal sulcus (IPS)(17–22). While these regions serve as the physiological basis for stereoscopic depth perception, the underlying neural activity covaries with disparity magnitudes within detectable ranges(23) across both visual streams. Preston TJ and colleagues(24) found a positive correlation of BOLD signal with the disparity magnitude for dorsal visual stream while having no correlation with the ventral visual stream. Subsequently, Wang F and colleagues(25) utilized a larger range of binocular disparities to investigate this disparity-response curve and confirmed the dominance of the dorsal visual stream; but, they did not find a monotonically increasing functional magnetic resonance imaging (fMRI) response with the disparity magnitude. However, the observed correlation in these studies is not intrinsic but instead arises as a result of specific causal interactions within their integrated network(26). Understanding these causal interactions would provide valuable insights into the neural underpinnings of the variations in stereoacuity thresholds observed among different individuals. For example, higher stereoacuity thresholds have been reported in patients with impaired eye alignment disorders such as amblyopia(7), strabismus(27) and induced anisometropic populations(28). Moreover, visually healthy controls could also exhibit sub-normal stereoacuity. Deepa and colleagues(29) reported that only around 13% of their tested population met the criterion for the normal level of stereopsis. Almost 45% of their study population had borderline stereopsis and the remaining 42% had reduced stereopsis. If the measuring test is kept the same, then perceptually this can only happen if there is distinct neural processing of varying disparity sizes. Therefore, changes in effective connectivity can serve as important biomarkers for extracting clinically relevant information from patients with impaired stereopsis. Recently, one study(30) utilized resting state fMRI to investigate causal interactions among several cortical regions in amblyopic patients. The authors report that the stereoscopic anomalies present in the amblyopic patients may result from the changes in effective connectivity of the higher-order visual regions. However, the use of resting-state functional MRI (rs-fMRI) as an experimental design would limit the interpretations of relationship between the topology of brain networks and the stereoscopic depth perception or to other aspects of brain function(31,32). Thus, it remains an open question how the complex topological properties of human brain networks are related to adapting to changing disparity magnitudes. This holds significant importance towards developing a holistic understanding of stereoscopic depth perception.

Our analysis is based on Granger Causality (GC)(33) – a widely used approach in exploring brain network causality – functional segregation (within-network connectivity) and functional integration (between-network connectivity)(34,35). Specifically, we use GC to construct directed networks to derive degree (D) and participation coefficients (PC)(36). Each node in the community structure has a distinct role depending upon their D and PC(36,37). The high-degree nodes that tend to be closely connected among their communities are called the *rich clubs*(38), whereas the higher PC nodes are called diverse clubs(39). Until recently, the rich club was thought to be critical for global communication and considered as an integrative and stable core of brain regions that coordinates the transmission of information across the network(38,40). However, Bertolero MA and colleagues(39) suggested that the higher PC nodes tend to interact even more strongly with other communities. While the brain network changes under different experimental conditions(41–43), a comprehensive understanding of rich and diverse clubs can help identify the neural substrate of disparity magnitude processing. Further, in order to investigate the existence of rich-diverse dichotomy under a clinically relevant range of disparity magnitudes, we appropriately modify our digital version (Digital Stereoacuity Test – DST(44)) of a random dot-based test – TNO (The Netherlands Organization) and subsequently utilize it within the fMRI setting. We chose TNO because of its superior performance to contour-based stereotests(45,46). Although TNO is one of the most widely used clinical tests for measuring stereopsis, employing well-controlled and clinically comparable stereoscopic stimuli in an fMRI setting can further help aid clinicians in establishing a stronger correlation between TNO thresholds and the fMRI results of the patients.

Another obvious related question concerns the interhemispheric differences associated with disparity processing. Some studies(23,47,48) have suggested bilateral and right hemispherically inclined roles of areas V3A and IPS, respectively, for the extraction and processing of stereoscopic depth perception. Contrarily, Wang F and colleagues(25), reported the involvement of V3, V3A and MT+ only in the right hemisphere. Moreover, other studies(19,49) have also identified a general dominance of the right hemisphere in the perceptual processing of stereopsis. These studies provide mixed evidence regarding the processing of disparity across hemispheres. Interestingly, there have been no studies addressing the interhemispheric differences specific to the processing of disparity magnitudes.

In this study, we hypothesized that: 1) there are unique rich and diverse clubs catering to processing of different disparity magnitudes, 2) if the identified rich and/or diverse club members yield a greater disparity-decoding performance and rank higher in feature importance, they are deemed to play significant roles in the processing of disparity magnitudes. 3) there are interhemispheric differences in the significant rich and/or diverse club members across different disparity magnitudes. Overall, our hypotheses revolve around the idea that the rich and diverse clubs play distinct roles in the complex brain topology and further contribute to revealing unique and distinct patterns during stereoscopic depth perception.

## 2. Material and methods

### 2.1. Participants

Twenty-six visually healthy controls (26 males; mean age: 26.81±3.22 years) with BCVA of 6/9 (0.67 or ≤ 0.17 logMAR) or better in both eyes participated in the Functional Magnetic Resonance Imaging (fMRI) based static-depth experiment. The data from 6 participants were discarded due to excessive head movement and/or inability to follow instructions during the experiment. Thus, all analyses were based on the data of remaining 20 participants (mean age: 26.15±3.18 years). All participants were chosen with normal stereoacuity (60 arc-sec) measured with the TNO stereo test. All participants were required to answer a post-experiment questionnaire (Supplementary material Appendix: 1) designed to understand qualitative aspects of their performance during the experiment. Participants were well informed of all experiments performed in this study and gave written consent prior to their participation. Participant recruitment and conduct of experiments were approved by ethics committee of All India Institute of Medical Sciences, New Delhi, India (IEC-511/17.06.22). This study was in accordance with the tenets of the Declaration of Helsinki.

### 2.2. MRI acquisition

Functional MRI data were acquired using a 3.0 T GE Scanner (Discovery MR 750w) equipped with a 32-channel phased-array head coil. The scanning parameters were as follows: TR = 2000 ms, TE = 29 ms, 176 slices, voxel resolution = 3 mm × 3 mm × 3.5 mm, slice thickness= 3 mm, Spacing: 0.5, FOV = 192 mm× 192 mm, flip angle = 65°, 610 volumes. The duration of the BOLD scan was 20 minutes. Subsequently, a T1-weighted anatomical image (TR = 15 ms, TE = 6.68 ms, voxel resolution = 0.5 mm × 0.5 mm × 1 mm, FOV = 176 x 176 mm, Flip angle = 10°) was acquired for 5 minutes. Foam pads were used to reduce scanning noise and minimize head motion.

### 2.3. Stimuli and procedure

We adapted the stimuli from our previously clinically validated stimuli(44) to align with the display specifications in the fMRI setting. The experiment consisted of a random dot stereogram (RDS) stimulus which was designed using Psychtoolbox v.3.0.17 and MATLAB R2020b. The stereoscopic stimuli differed from our previous DST design in two aspects: (i) The RDS square contained a hidden 3D shape (‘⊔’ or ‘⊓’ of disparities 120, 240, 480 & 800 arc-secs) where the ‘⊔’ shape appeared during 800 and 240 arc-sec conditions and ‘⊓’ appeared during 480 and 120 arc-sec conditions and, (ii) the size of dots and RDS square were appropriately scaled to fit in the Nordic Neuro Lab (NNL; NordicNeurolab, Bergen, Norway) Visual System goggles (Resolution: 800×600, Refresh rate: 85 Hz, FOV: 28.6° horizontal x 20.3° vertical) used inside MRI scanner. The red and blue anaglyph filters were superimposed over the left and right eye lenses, respectively. The luminance of the dots as seen through the red lens was 4.6 and 4.3 cd/m^2^ as seen through the blue lens. The stimuli were presented in a blocked-design including 3D-shape (‘⊔’ or ‘⊓’), 2D scrambled-noise (SN) (RDS square without hidden 3D shape) and fixation (+) blocks. Each block was repeated five times except the fixation block, which repeated after every 2 experimental (3D-shape and 2D SN) blocks. Each block lasted 20 seconds with a total experiment duration of 20 minutes (excluding 5 minutes of T1-structural scan). Participants had to indicate the shape hidden in the RDS square with a button press (Lumina 3G controller) inside the scanner (left for ‘⊔’ and right for ‘⊓’). Each participant was subjected to a 1-hour task-training session with NNL goggles prior to the start of the actual experiment. This was done to improve the perceptual learning of stereopsis(50) and keep a fair comparison among the participants. Fig. 1 illustrates the overall experimental setup and the fMRI block design used in the static depth experiment.

**Fig. 1.**
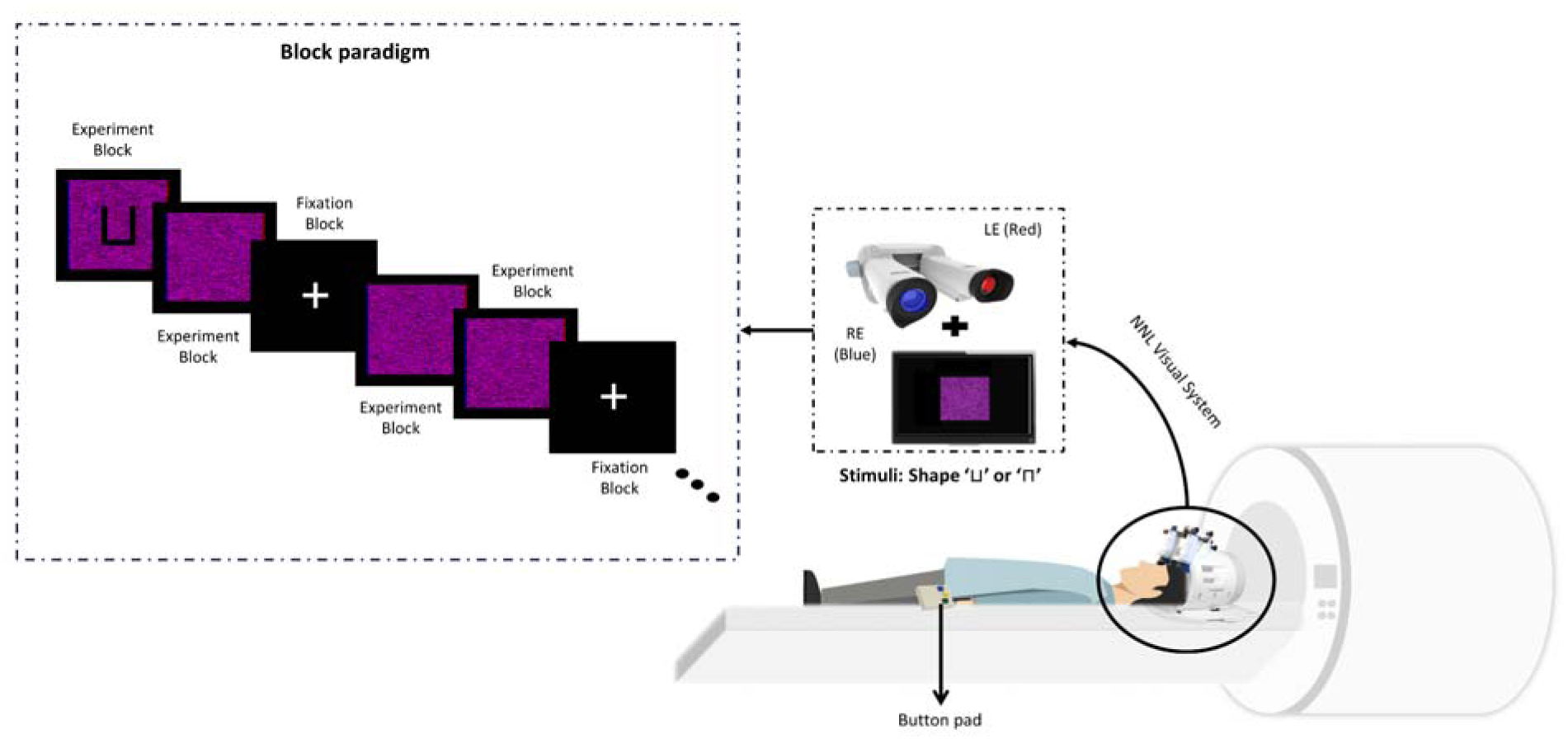
Participants lay in supine position in the MR scanner with their left- and right-hand index fingers placed on separate button pads (Lumina 3G controller) to indicate the 3D-shape (‘⊔’ or ‘⊓’) in the static depth experiment. The fMRI block design (left inset view) shows the occurrence of two experimental blocks followed by a fixation block wherein each block lasted for 20 seconds. The ‘⊔’ shape in the experimental block is for illustration purposes only. The actual ‘⊔’ or ‘⊓’ shape was visible only when viewed stereoscopically with the NNL visual goggles (right inset view) and the superimposed red and blue films on left and right eye, respectively.

### 2.4. fMRI data analysis

Functional MRI data were minimally preprocessed using fMRIPrep-21.0.1(51) based on Nipype 1.5.127(52) and the structural T1 data were preprocessed using FreeSurfer (version: 6.0.0)(53). The Analysis of Functional Neuroimages (AFNI) programs *3dmerge* (with full width half maximum of 4mm), *3dcalc* and *3ddtrend* were used to spatially smooth, scale and detrend fMRI data, respectively. Next, *3dDeconvolve* was used to perform first-level general linear model analysis to extract voxel-wise response amplitude for 3D disparity conditions, 2D SN and fixation blocks. To control the false positive rate (FPR), AFNI program *3dttest++* was used with the *Clustsim* option for randomization and permutation simulations to produce cluster-level threshold values. Clusters were defined as groups of voxels above the uncorrected significance threshold whose faces or edges touched as the default setting of AFNI. This revealed the minimum size of a voxel cluster needed for a corrected *p* of 0.001. Subsequently, conjunction analysis(54) was performed to obtain the significant activation clusters across all disparity magnitudes. A region of interest (ROI) of size 5 mm was created at the peak of activation cluster using AFNI programs *3dUndump* and *3dfractionize* across each hemisphere. Finally, the ROIs were defined based on the Glasser HCP 2016 surface-based parcellation atlas(55).

### 2.5 Network Construction using Granger Causality

Granger Causality (GC) was employed using multivariate auto-regressive modelling (MAR)(56) with model and lag selection based on Akaike Information Criterion (AIC) using *1dGC.R*(56) on the ROI-time series extracted from the detrended fMRI data. Subject motion parameters were considered confounders in the MAR model. The resulting weighted path matrices from *1dGC.R* consisted of only statistically significant connections at p<0.05.

### 2.6 Definition of rich and diverse clubs

The weighted path matrices were randomized with positively and negatively signed connections while preserving the positively and negatively signed in-degree (D_in_) and out-degree (D_out_) distributions(57). We computed node-wise D_in_, D_out_ participation-in (P_in_) and participation-out (P_out_) coefficients to observe inter-regional differences across disparity magnitudes using the Brain Connectivity Toolbox (BCT)(58) wherein, in-and out refer to the incoming and outgoing connections, respectively. Next, we defined the brain regions with high (median-value over 45^th^ percentile and above) average degree (D_avg_, wherein average is over in-and-out connections) as the *rich club* members across each disparity condition(39). This follows the notion that nodes with higher degrees are inclined to connect with each other intensely(38). We reduced the percentile cut-off from the 80^th^ to 45^th^ percentile to account for fewer nodes in our network. While no analogous diverse club coefficient is specified in the literature, we utilized the same threshold for the diverse club members. Subsequently, the brain regions with the high (median-value over 45^th^ percentile and above) average participation coefficient (PC_avg_, wherein average is over in-and-out connections) were defined as *diverse club* members. A region was termed as *overlapping region* if it was present in both rich and diverse categories and was considered to have the highest importance across all regions because of its higher D and PC(59). We took the same ROIs across each hemisphere in the GC analysis for a matched comparison. Finally, the Kruskal-Wallis test (at *p*<0.05 & Bonferroni-corrected) was used to highlight significant global and local features of these rich and diverse regions across each hemisphere. This test was used as an alternative to its parametric equivalent – one-way analysis of variance (ANOVA).

### 2.7 Decoding disparity magnitudes across right and left hemisphere

To further validate the importance of the identified rich and diverse club members in terms of their ability to decode disparity magnitudes, we trained a decision tree (DT) model(60) based on the 3 disparity conditions (800 arc-sec, 240 arc-sec and 480 arc-sec). DT models are white-box models known for their easy interpretability and ability to provide feature importance scores. We optimized the hyperparameters (impurity criterion, maximum depth of tree, minimum samples in the split and the type of splitter) using a 5-fold grid search cross-validation. We performed this analysis for RH and LH separately to elucidate the interhemispheric differences in decoding disparity magnitude.

## 3. Results

### 3.1 Group-level fMRI activations across different disparity magnitudes

The psychometric curve (Fig. 2) showed an overall increase in percent correct responses with disparity magnitude except at 480 arc-sec wherein the performance was greater compared to 800 arc-sec condition. The 120 arc-sec condition showed a less than chance (Pc: Proportion correct=0.33) performance and was therefore excluded in further analyses. Next, we perform group analysis followed by cluster correction (using *Clustsim* described in Section 2.4) which revealed activation maps for contrasts 800 > SN (Fig. 3a), 480 > SN noise (Fig. 3b) and 240 > SN (Fig. 3c). The clusters that survived the uncorrected cluster-forming *p*-threshold of p<0.001 for overall alpha (probability that the given cluster is greater than cluster-size threshold) threshold of p<0.05 for all three conditions are detailed in Supplementary Table 1. We also observed consistent activations in post-central and pre-frontal cortex pertaining to button-press(61) and generic task executions(62,63) in all disparity-task conditions. Since these were not the primary focus of our research hypotheses, we only selected the clusters associated with stereoscopic depth perception task. Further, Supplementary Table 2 shows the details of all ROIs derived from these clusters across both hemispheres.

**Fig. 2.**
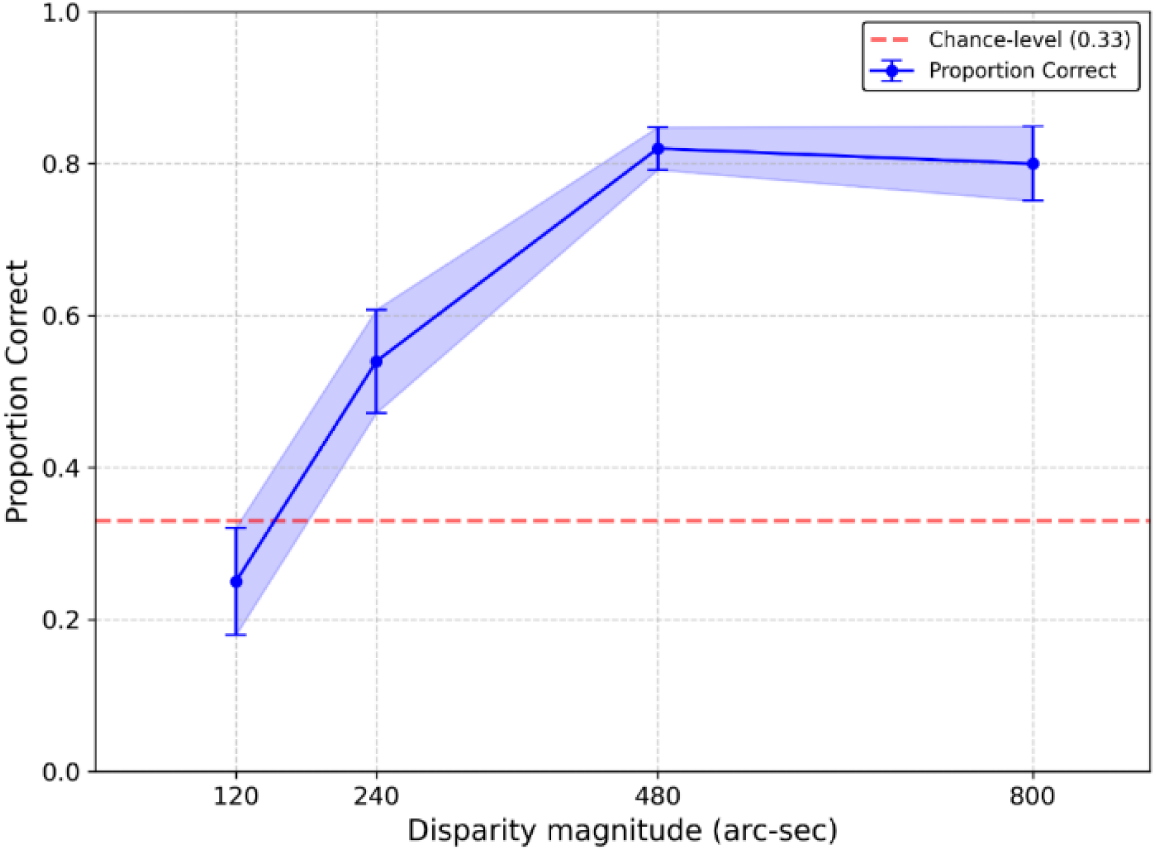
The psychometric curve showing increase in performance of participants (N=20) with th disparity magnitude under the fMRI stereoscopic depth perception task.

**Fig. 3:**
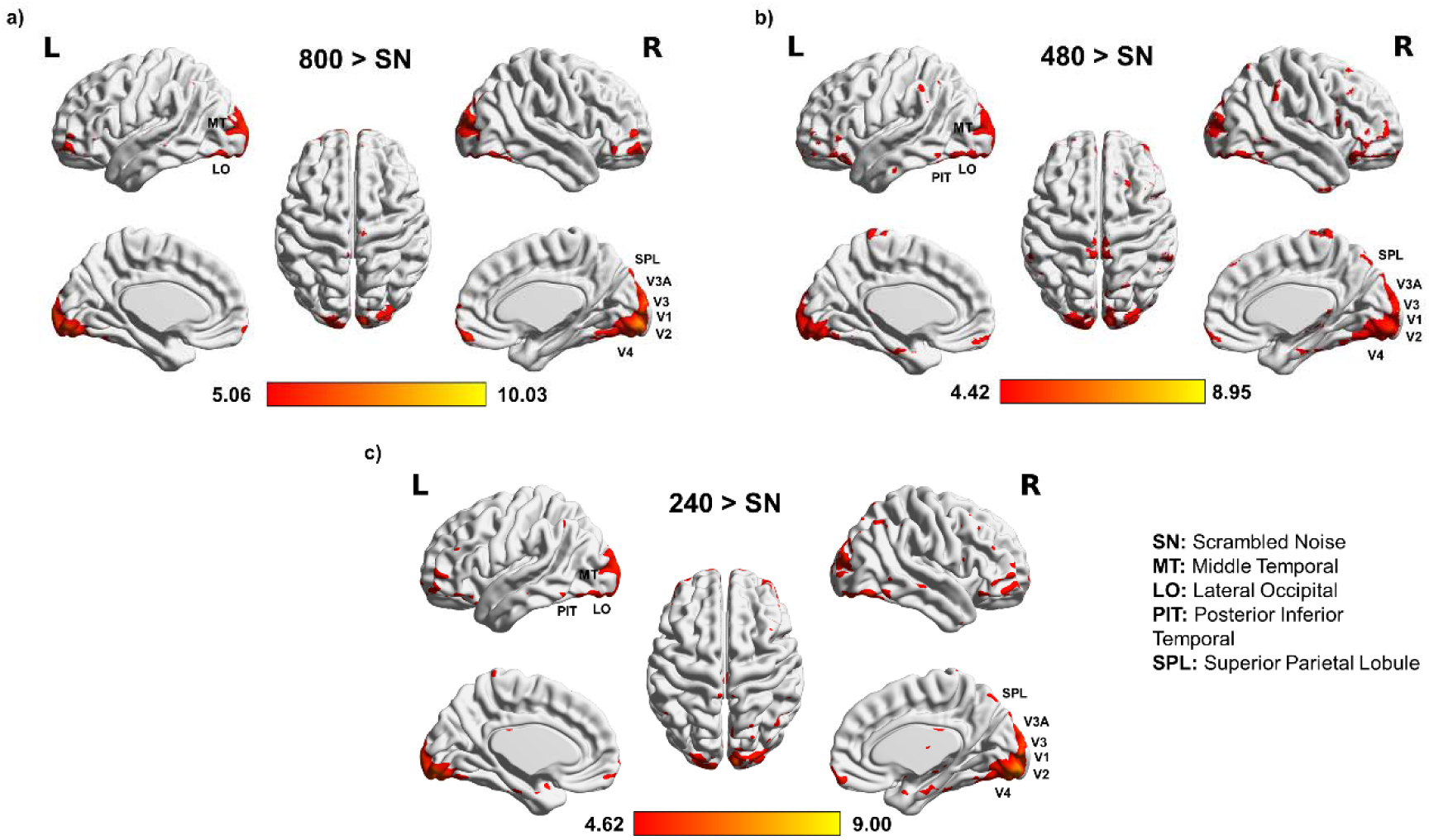
(a) Group fMRI activation patterns (N=20) for the GLM contrast [800 > SN] at the cluster defining threshold of p = 0.001 (cluster > 39 voxels for it to be significant at p<0.05), (b) Group fMRI activation patterns (N=20) for the GLM contrast [480 > SN] at the cluster defining threshold of p = 0.001 (cluster > 32 voxels for it to be significant at p<0.05), (c) Group fMRI activation patterns (N=20) for the GLM contrast [240 > SN] at the cluster defining threshold of p = 0.001 (cluster > 29 voxels for it to be significant at p<0.05). The details of activation clusters are mentioned in supplementary Tables 1&2.

### 3.2 The rich and diverse clubs across the right hemisphere

In the next step, we constructed networks by applying GC to the ROIs derived (Supplementary Table 2) from activation analysis. The D_avg_ and PC_avg_ obtained using BCT across all regions and disparity conditions for RH are shown in supplementary figures S1-S6. Next we selected the brain regions with high D_avg_ (median-value over 45^th^ percentile and above) and found that the rich club members included: 1) MT, V2*, SPL*, LO^+^, V1^+^, V4^+^ and PIT^+^ during the 800 arc-sec condition where regions with * and ^+^ had equal D_avg_ respectively, 2) regions LO, MT*, SPL*, V1*, V2*, PIT* during the 240 arc-sec condition and 3) regions MT, SPL*, LO*, V1* and PIT* during the 480 arc-sec condition. By selecting the brain regions with high PC_avg_ (median-value over 45^th^ percentile and above), we found that the diverse club members included: 1) LO, MT, V1 and V2 during the 800 arc-sec condition, 2) regions MT, PIT, LO and V4 during the 240 arc-sec condition and 3) regions V1, V2, LO, MT and V3 during the 480 arc-sec condition. The common rich and diverse club members included: 1) MT, LO, V1 and V2 during the 800 arc-sec condition, 2) regions MT, LO, and PIT during the 240 arc-sec condition and 3) regions MT, V1 and LO during the 480 arc-sec condition.

For further analysis, we used the Kruskal-Wallis test coupled with post-hoc tests (Tukey-Kramer) for pairwise comparisons across 3D conditions that showed: 1) D_in_ of MT was significantly different (χ^2^=8.95, df=2; p=0.011) for the 240 and 480 arc-sec conditions (median: 3 & 2; p=0.0135; see Fig. 4a) respectively, 2) D_out_ of regions V2 (χ^2^=6.52, df=2; p=0.038) and SPL (χ^2^=6.24, df=2; p=0.044) were significantly different for the 240 and 480 arc-sec conditions (median: 5 & 2; p=0.036; see Fig. 4b and median: 4 & 2; p=0.033; see Fig. 4c for V2 and SPL respectively), 3) P_in_ coefficient of V2 was significantly different (χ^2^=6.29, df=2; p=0.042) for the 240 and 480 arc-sec conditions (median: 0 & 1.248; p=0.0324; see Fig. 4d) respectively, and 4) P_in_ coefficient of V3 (χ^2^=6.46, df=2; p=0.039) was significantly different for the 800 and 480 arc-sec conditions (median: 0 & 1.13; p=0.0439; see Fig. 4e) respectively.

**Fig. 4:**
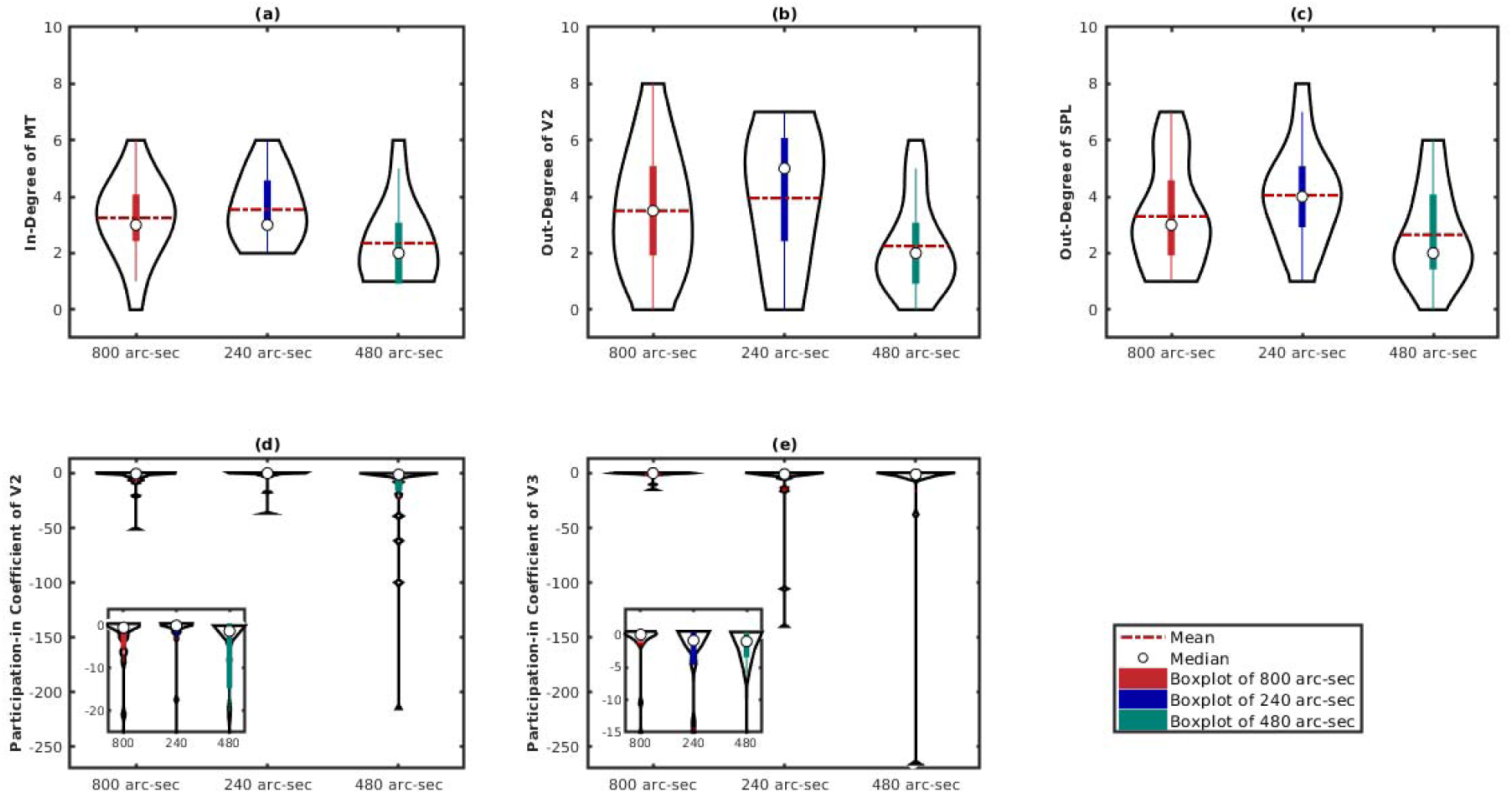
(a) In-Degree of region MT is significantly different for the 240 and 480 arc-sec conditions (median: 3 & 2; p=0.0135), (b) Out-Degree of region V2 is significantly different for the 240 and 480 arc-sec conditions (median: 5 & 2; p=0.036), (c) Out-Degree of region SPL (median: 4 & 2; p=0.033) is significantly different for the 240 and 480 arc-sec conditions, (d) Participation-in coefficient of region V2 is significantly different for the 240 and 480 arc-sec conditions (median: 0 & −1.248; p=0.0324), (e) Participation-in coefficient of V3 is significantly different for the 800 and 480 arc-sec conditions (median: 0 & 1.13; p=0.0439). The inset figures illustrate the zoomed view of the corresponding subplots.

### 3.3 The rich and diverse clubs across the left hemisphere

We follow a similar analysis for LH as we did for RH. The D_avg_ and PC_avg_ are illustrated in in supplementary figures S7-S12. The brain regions included in the rich club included: 1) V2, MT, V4 and PIT during 800 arc-sec condition, 2) regions MT*, V2*, PIT*, V1^+^, V3A^+^ and SPL^+^ during the 240 arc-sec condition and 3) regions LO, V4, MT*, V1*, PIT*, V3A* and SPL* during the 480 arc-sec condition. Meanwhile, the diverse club included: 1) regions SPL, V2, MT and V1 during the 800 arc-sec condition, 2) regions PIT, MT, V2 and V3A during the 240 arc-sec condition and 3) regions LO, V1, PIT and MT during the 480 arc-sec condition. The common rich and diverse club regions included: 1) V2 and MT during the 800 arc-sec condition, 2) MT, PIT and V2 during the 240 arc-sec condition and 3) LO, PIT and MT during the 480 arc-sec condition. The Kruskal-Wallis test coupled with post-hoc tests for pairwise comparisons across 3D conditions revealed that the P_out_ coefficient of V2 is significantly different (χ^2^=7.98, df=2; p=0.018) for the 800 (p=0.039) and the 480 (p=0.037) arc-sec conditions as compared to the 240 arc-sec condition (median: 0.3716, 0.316 & 0 respectively; see Fig. 5).

**Fig. 5:**
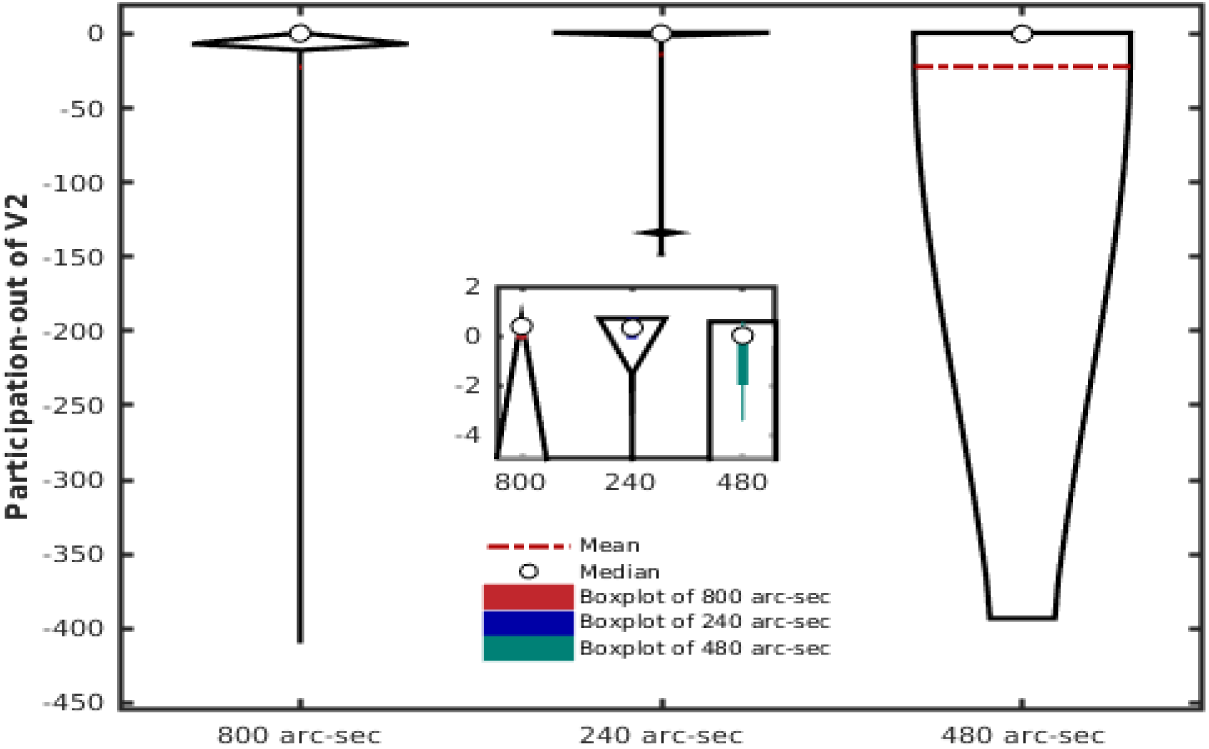
Participation-out coefficient of V2 (median: 0.3716, 0.316 & 0) is significantly different for 800 (p=0.039) and 240 (p=0.037) arc-sec as compared to 480 arc-sec condition. The red line in the middle of each box represents the median. The inset figure illustrates the zoomed view of the original plot.

### 3.4 Decoding disparity magnitude with diverse club members across right hemisphere

The input features of DT consisted of both P_in_ and P_out_ coefficients of all ROIs. Fig. 6(b) shows the DT model that was trained to classify disparity conditions based on the above-mentioned features. The model identified the best parameters (criterion: entropy, max depth: 4, min sample: 2 and splitter: random) using a 5-fold grid search cross validation to optimize the hyperparameters and was able to classify with precision: 0.967, recall: 0.945, and F1 score: 0.959. The weight assigned to each feature (i.e., feature importance) by the decision tree algorithm is mentioned in Fig. 6(a).

**Fig. 6:**
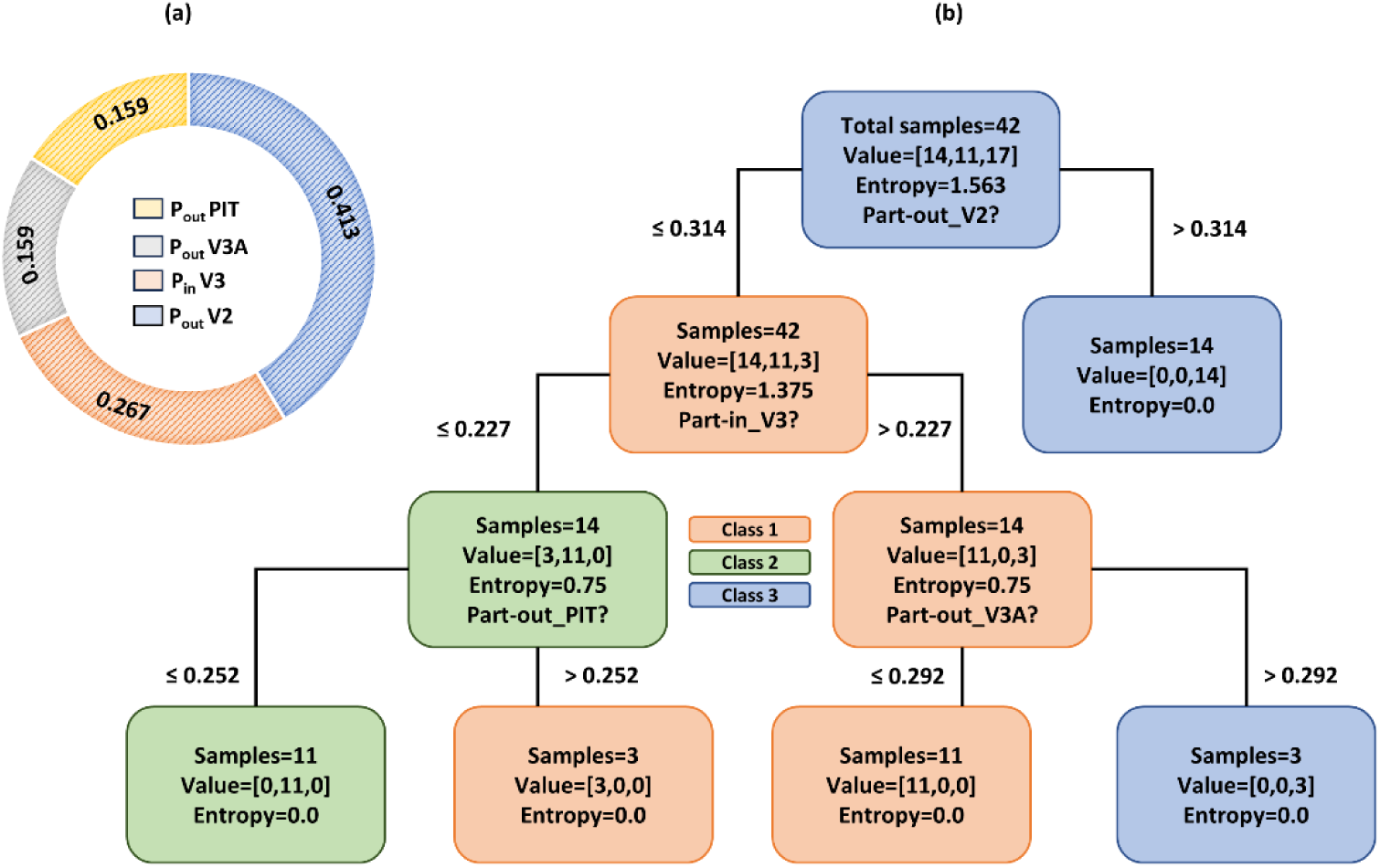
(a) Feature importance evaluated by the Decision tree model for decoding disparity magnitudes in the Right hemisphere, (b) Diverse club decision tree model for classifying disparity conditions – 800 arc-sec (class 1), 240 arc-sec (class 2) and 480 arc-sec (class 3) across right hemisphere. The value set at each level of the tree indicates the number of correctly classified and misclassified disparity conditions.

### 3.5 Decoding disparity magnitude with rich club members across right hemisphere

Unlike the P_in_ and P_out_ coefficients across each node, the D_in_ and D_out_ features did not result in accurate classification. The performance of the decision tree model was merely at chance level (33%) with precision: 0.357, recall: 0.296, and F1 score: 0.273. Consequently, to investigate whether this low performance was due to the higher similarity in the processing of any two disparity magnitudes or the poor decoding of disparity magnitudes by rich clubs, we trained the decision tree classifier pairwise. The model performance, however, above chance level (>33%), was still worse when compared to diverse clubs in all three pairwise conditions – class 1 vs class 2 (precision: 0.583, recall: 0.586, and F1 score: 0.584), class 2 vs class 3 (precision: 0.413, recall: 0.416, and F1 score: 0.415) and class 1 vs class 3 (precision: 0.444, recall: 0.452, and F1 score: 0.436).

### 3.6 Decoding disparity magnitude with diverse club members across left hemisphere

We followed a similar approach for LH as described in section 3.4 and trained our model with P_in_ and P_out_ coefficients of all ROIs. The decision tree algorithm (shown in Fig. 7b) could decode the disparity conditions with precision: 0.834, recall: 0.857 and F1 score: 0.802. The weight assigned to each feature by the decision tree algorithm is mentioned in Fig. 7(a).

**Fig. 7:**
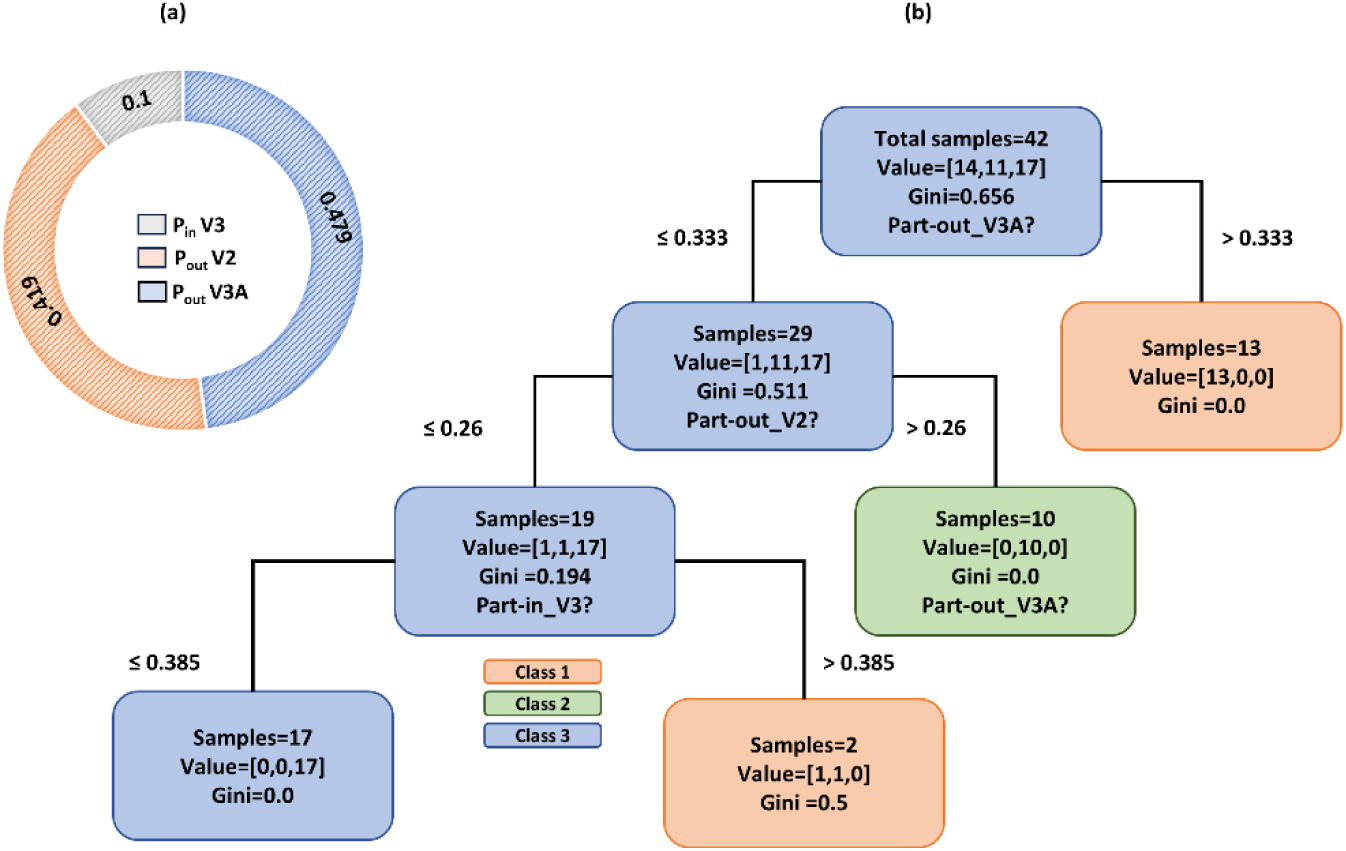
(a) Feature importance evaluated by the Decision tree model for decoding disparity magnitudes in the Left hemisphere, (b) Diverse club decision tree model for classifying disparity conditions – 800 arc-sec (class 1), 240 arc-sec (class 2) and 480 arc-sec (class 3) across left hemisphere. The value set at each level of the tree indicates the number of correctly classified and misclassified disparity conditions.

### 3.7 Decoding disparity magnitude with rich club members across left hemisphere

The decision tree model was trained with D_in_ and D_out_ features for all ROIs. As expected, the model performed similarly to the right hemisphere for the rich clubs with precision: 0.424, recall: 0.404, and F1 score: 0.366.

### 3.8 Differences between right and left hemisphere

Kruskal-Wallis test revealed that PC_avg_ was significantly different across several brain regions in LH and RH. The box plot visualization of each significant finding is shown in S13 to S21 and the corresponding statistical values are mentioned Table 1. There were no other significant inter-hemispheric differences among the remaining conditions.

**Table 1.** Interhemispheric differences in the average participation coefficient across different disparity magnitudes. The differences are significant at p<0.05.

| SNo. | ROI | $\chi^2$ & df | *p* value | Disparity Magnitude (arc-sec) | Median: RH & LH |
| --- | --- | --- | --- | --- | --- |
| 1 | V1 | 5.57 & 1 | 0.018 | 240 | 0.376 & 0.23 |
| 2 | V1 | 6.26 & 1 | 0.0123 | 480 | 0.307 & 0.283 |
| 3 | V2 | 5.94 & 1 | 0.014 | 240 | 0.333 & 0.22 |
| 4 | PIT | 7.057 & 1 | 0.0079 | 800 | 0.353 & 0.325 |
| 5 | PIT | 8.23 & 1 | 0.0041 | 240 | 0.387 & 0.227 |
| 6 | V3A | 11.49 & 1 | 0.000696 | 240 | 0.426 & 0.225 |
| 7 | V3A | 11.86 & 1 | 0.000572 | 480 | 0.28 & 0.356 |
| 8 | SPL | 15.18 & 1 | 0.0000973 | 240 | 0.341 & 0.251 |
| 9 | SPL | 6.269 & 1 | 0.0124 | 480 | 0.353 & 0.321 |

## 4. Discussion

The main findings of our study indicate that the effective connectivity between brain regions changes with the disparity magnitude for visually healthy controls. Specifically, our results reveal the presence of distinct rich and diverse clubs that vary across different disparity magnitudes. Furthermore, our analysis indicates that the MT region serves as the only common rich and diverse region across all disparity magnitudes and in both hemispheres. We also find that diverse clubs exhibit better performance in decoding disparity magnitudes, thereby providing further support to the growing evidence that diverse clubs are indeed the integrative core for processing disparities. Finally, we find that subtle inter-hemispheric differences exist across disparity conditions. Below, we discuss these findings in more detail.

### 4.1 Distinct rich and diverse clubs exist under different disparity magnitudes

The pairwise comparisons of D_in_, D_out_ and P_in_, P_out_ (representing rich and diverse club nature, respectively) of all ROIs from Kruskal Wallis analysis revealed significant differences under different disparity magnitudes (Fig. 4). Specifically for RH, stereoscopic viewing in the 240 arc-sec condition resulted in a higher D_in_ for MT, and D_out_ for V2 and SPL as compared to 480 arc-sec condition (Figures 4a-c). Similarly, participants viewing the 480 arc-sec condition exhibited a significantly higher P_in_ in regions V2 and V3 compared to both 240 and 800 arc-sec conditions, respectively (see Figures. 4d-e). The fluctuations in D and PC observed over the range of disparities are expected as the fMRI response itself does not have a strictly monotonic relationship with disparity magnitude(23–25,64). Backus and colleagues(23) found that mean response in early visual areas is a bimodal function of disparity magnitude, where BOLD activity first increases from 30 to 60 arc-sec, then decreases until 225 arc-sec, and finally rises to 900 arc-sec. While their findings support correlation between BOLD activity and disparity magnitudes (similar to our experiment), we argue that the existence of significantly different rich and diverse clubs may explain distinct neural mechanisms beyond the interpretation of BOLD activity changes alone.

In the subsequent subsections, we explain the region-wise contributions in decoding disparity magnitudes obtained from DT analysis. We focus only on the RH due to its higher DT-classification performance compared to LH.

#### 4.1.1 Common stereo processing until V2

Despite having a significantly higher P_in_ in region V2 for 480 arc-sec condition compared to both 240 arc-sec conditions, the DT classification pathway starts with a common region (Fig. 6b) V2 for all disparity magnitudes. This may be because early visual areas provide crude yet important representations of disparity(20,65) to the extent that lesions to them can lead to impaired stereoacuity(66–68). Moreover, the shared utilization of V2 across all disparity magnitudes could also be reasoned with the “correspondence problem” in stereopsis. For the visual system to match features in the left eye with the corresponding features in the right eye, it must reject a large number of possible local matches between the images of the two eyes while preserving the correct matches. This entire corresponding feature matching across eyes takes place in the extrastriate cortex(69), with specialization for relative disparity processing in V2(70), irrespective of the disparity magnitude.

#### 4.1.2 Role of V3A and PIT in decoding disparity magnitudes

In our experiment, V3A discriminated between 800 (coarse) and 480 arc-sec (mid) disparity magnitudes. There could be three reasons for this: 1) V3A neurons are highly sensitive(18,23,48,71,72) towards discriminating disparity magnitudes in general, 2) V3A specifically aids in encoding mid-level disparities. However, the range of disparity categorized as coarse, mid or relatively fine is subject to its definition by the experimenter. For instance – Chen and colleagues(73) used disparities less than 600 arc-sec in their “smallest detectable disparity” experiment and concluded the role of V3A in decoding the finer disparity signals. Although we refer to 480 arc-sec as a mid-range disparity, our finding is in line with Chen and colleagues(73), 3) V3A is exclusively a diverse club member during the 480 arc-sec condition defined by a higher PC (> 0.292 for 480 arc-sec when compared with < 0.292 for 800 arc-sec) that in turn aided the decoding.

Furthermore, PIT effectively distinguishes between 800 (coarse) and 240 arc-sec (relatively finer) disparity magnitudes (Fig. 6b). Notably, as these conditions represent the largest and smallest disparities in our analysis, PIT neurons likely exhibit sensitivity in discriminating between both ranges. While limited evidence exists in humans(74), non-human primate studies link PIT to fine(75,76) and relatively coarse depth discrimination tasks(77). However, PIT’s role extends beyond disparity processing, playing a significant part in decoding 3D shape from stereopsis(68,76,78–80). Interestingly, PIT also served as a common rich club across all disparity magnitudes (see section 3.2), potentially contributing to the shared processing of 3D shape from stereopsis. Further research on PIT’s specific role as a rich or diverse club in decoding shape and magnitude is essential to corroborate our results.

### 4.2 The rich and diverse nature of MT

MT exhibited remarkable overlapping rich and diverse properties across disparity magnitudes and both hemispheres. While MT is known to have a selective preference for disparity processing(81–84), intriguingly, the decision tree algorithm did not reveal MT as a prominent feature in decoding disparity magnitudes. This finding could be attributed to MT’s consistent contribution across all disparity magnitudes, encompassing coarse, mid, and relatively finer ranges. Notably, Uka, T., & DeAngelis, G. C(85) and Neri, P and colleagues(47) argue MT to be a part of the neural substrate underlying only coarser disparities. Contrarily, Krug, K., & Parker, A. J(86) utilized a wide range of disparities (0 to 4320 arc-sec) and summarized that V5/MT neurons are also selective for relative disparity. However, their distinction of fine and coarse disparities is based on the type of task employed – absolute (coarse) vs relative (fine) – rather than the magnitude of test disparities itself.

Although our experiment is designed to test disparity magnitudes with only relative disparities, our findings emphasize that the dual – rich and diverse – characteristic of MT imply its possible contribution to the primitive stereo processing required for all disparity magnitudes rather than being exclusively dedicated to coarser or finer disparities.

### 4.3 Diverse clubs than rich clubs are indeed integrative core in stereo processing

The superior performance of diverse clubs (see section 3.4) in predicting disparity magnitudes across both hemispheres provides compelling evidence that PC is a better indicator of the underlying neural mechanisms of disparity magnitudes as opposed to D, which merely measures node connections within or between networks. Our findings align with the recent evidence(39) which indicates that diverse clubs exhibit characteristics of an integrative network function to a larger extent compared to rich clubs. Besides, the rich clubs primarily serve to facilitate the formation of specialized subnetworks(38). While rich and diverse clubs may play distinct roles in the brain network communication(59), intuitively, this points to a possibility that the rich clubs are more involved in functional segregation than integration. Considering the global nature of functional integration and its association with diverse clubs, this may be a plausible basis for their superior performance over rich clubs in decoding disparity magnitudes.

### 4.4 Inter-hemispheric differences across disparity magnitudes

Table 1 highlights the significant inter-hemispheric differences in the average PC across V1, V2, PIT, V3A and SPL. Conforming to the previous studies(19,49,87), we found an overall larger involvement of RH in the perception of stereopsis across regions V1, V2, PIT and SPL. This is in line with our preliminary findings(88) with fewer participants (n=11) where we found that SPL facilitates functional integration for the mid-level disparity magnitude. Besides, SPL is known to exhibit a right hemispheric bias(19,48,89,90) in the disparity processing. Contrary to SPL, the interpretation of hemispheric dominance in V3A is unclear with previous studies suggesting a bilateral(18,23,47,91) and right hemispheric dominance(25). In our study, we found RH and LH dominance during 240 arc-sec and 480 arc-sec conditions respectively. This leads to the possible dependence of hemispheric asymmetry of V3A on the magnitude of disparity. Therefore, except for V3A, we suggest an overall dominance of RH in stereo processing at least for the regions analyzed in our study.

### 4.5 The advantage of a stereoscopic task-based effective connective study over resting-state studies

Overall, our study builds upon previous research highlighting aberrant functional connectivity in the primary(92) and higher-order visual cortex(93). The only study that we can compare our results to is that by Chen and colleagues(30). They used Dynamic Causal Modelling from resting state fMRI to highlight connectivity abnormalities (V2 to LO) in amblyopic patients. They report abnormal effective connectivity specifically in V3d, V3A, V3B, and LO regions (they state them as “important nodes” in the network) by comparing the networks for amblyopic patients and visually healthy controls. Our results are consistent in terms of node-wise importance during stereoscopic depth perception. However, it is important to note that their findings stem from topological comparison between the two groups in the absence of a stereoscopic depth perception task. Therefore, their results represent the intrinsic states of amblyopic visual system and restricts the extent to which their findings can be meaningfully interpreted. On the other hand, our study is specific and designed for a deeper understanding of the neural mechanisms under various disparity magnitudes in a controlled stereoscopic depth perception task. These advantages contribute to a nuanced understanding of neural mechanisms behind stereoscopic depth perception. Moreover, the existence of distinct rich and diverse club patterns across different disparity magnitudes may explain the physiological basis for the variability in stereoacuities reported in visually healthy individuals(29). Our results provide complementary functional evidence to the structural evidence demonstrated by Oishi and colleagues(94). Notably, our findings also shed light on distinct rich and diverse club patterns across disparity magnitudes and may have potential implications for understanding eye misalignment disorders.

### 4.6 Limitation and future directions

There are some limitations to our present study. The NNL goggles are limited in terms of their resolution and FOV(95). This may have negatively impacted the participants’ performance during the 120 arc-sec disparity condition. Therefore, using some of the latest fMRI-compatible binocular devices with improved display capabilities(96) to investigate smaller disparities would be helpful. We also used anaglyph-based stereoscopic stimuli in our investigations. While this was by design so that we could validate the stimulus against the clinically used TNO depth test, future studies could make use of natural and ecologically valid 3D stimuli to confirm whether our findings extend to them.

## 5. Conclusion

Our investigations into the neural mechanisms of binocular disparity processing using Granger causality and graph measures revealed distinct rich and diverse clubs across different disparity magnitudes. Notably, area MT plays a specialized role with overlapping rich and diverse characteristics, and diverse clubs are preferred for decoding disparity magnitudes. While an overall right hemisphere dominance was observed in binocular disparity processing, subtle interhemispheric differences were found across various brain regions. Our study sets the stage for conducting further investigations on binocular disparity processing, particularly in the context of neuro-ophthalmic disorders with binocular impairments such as strabismus and amblyopia.

## Acknowledgments

This study received funding from the Department of Science and Technology - Cognitive Science Research Initiative Project #RP03962G, Govt. of India. The funding organizations had no role in the design, conduct or analysis of this research.

## Data Accessibility

The data that support the findings of this study are available from the corresponding author upon reasonable request.

## Additional Information

The preliminary results of the fMRI study in this manuscript have been presented as a poster at the Annual Meeting of the Organization for Human Brain Mapping, July 22–26, 2023, Montreal, Québec, Canada.

## References

1. Gonzalez F, Perez R. Neural mechanisms underlying stereoscopic vision. Prog Neurobiol. 1998;55(3):191–224.

2. Birch EE, Stager DR, Everett ME. Random dot stereoacuity following surgical correction of infantile esotropia. Vol. 32, Journal of Pediatric Ophthalmology \& Strabismus. SLACK Incorporated Thorofare, NJ; 1995. p. 231–5.

3. Birch EE, Fawcett S, Stager DR. Why does early surgical alignment improve stereoacuity outcomes in infantile esotropia? J AAPOS. 2000;4(1):10–4.

4. Fawcett S, Leffler J, Birch EE. Factors influencing stereoacuity in accommodative esotropia. J Am Assoc Pediatr Ophthalmol Strabismus. 2000;4(1):15–20.

5. Birch EE, Wang J. Stereoacuity outcomes after treatment of infantile and accommodative esotropia. Optom Vis Sci. 2009;86(6):647–52.

6. Mehner L, Ng SM, Singh J. Interventions for infantile esotropia. Cochrane Database Syst Rev. 2023;2023(1).

7. Levi DM, Knill DC, Bavelier D. Stereopsis and amblyopia: A mini-review HHS Public Access. Vis Res. 2012;114(2012):17–30.

8. Webber AL, Wood J. Amblyopia: Prevalence, natural history, functional effects and treatment. Clin Exp Optom. 2005;88(6):365–75.

9. Grant S, Melmoth DR, Morgan MJ, Finlay AL. Prehension deficits in amblyopia. Investig Ophthalmol Vis Sci. 2007;48(3):1139–48.

10. Niechwiej-Szwedo E, Kennedy SA, Colpa L, Chandrakumar M, Goltz HC, Wong AMF. Effects of induced monocular blur versus anisometropic amblyopia on saccades, reaching, and eye-hand coordination. Investig Ophthalmol Vis Sci. 2012;53(8):4354–62.

11. Suttle CM, Melmoth DR, Finlay AL, Sloper JJ, Grant S. Eye-hand coordination skills in children with and without amblyopia. Investig Ophthalmol Vis Sci. 2011;52(3):1851–64.

12. Wainman B, Pukas G, Wolak L, Mohanraj S, Lamb J, Norman GR. The Critical Role of Stereopsis in Virtual and Mixed Reality Learning Environments. Anat Sci Educ. 2020;13(3):401–12.

13. Levi DM. Applications and implications for extended reality to improve binocular vision and stereopsis. J Vis. 2023;23(1):14.

14. Poggio T, Torre V, Koch C. Computational vision and regularization theory. Nature. 1985;317(6035):314–9.

15. Pizlo Z. Perception viewed as an inverse problem. Vision Res. 2001;41(24):3145–61.

16. Nieder A. Stereoscopic vision: Solving the correspondence problem. Curr Biol. 2003;13(10):394–6.

17. Balasopoulou A, Κokkinos P, Pagoulatos D, Plotas P, Makri OE, Georgakopoulos CD, et al. Symposium Recent advances and challenges in the management of retinoblastoma Globe □ saving Treatments. BMC Ophthalmol [Internet]. 2017;17(1):1. Available from: http://www.ncbi.nlm.nih.gov/pubmed/28331284 http://www.pubmedcentral.nih.gov/articlerender.fcgi?artid=PMC5354527 http://bmcpsychiatry.biomedcentral.com/articles/10.1186/1471-244X-11-49 http://bmcophthalmol.biomedcentral.com/articles/10.1186/s12886

18. Goncalves NR, Ban H, Sánchez-Panchuelo RM, Francis ST, Schluppeck D, Welchman AE. 7 Tesla fMRI reveals systematic functional organization for binocular disparity in dorsal visual cortex. J Neurosci. 2015;35(7):3056–72.

19. Nishida Y, Hayashi O, Iwami T, Kimura M, Kani K, Ito R, et al. Stereopsis-processing regions in the human parieto-occipital cortex. Neuroreport. 2001;12(10):2259–63.

20. Roe AW, Parker AJ, Born RT, DeAngelis GC. Disparity channels in early vision. J Neurosci. 2007;27(44):11820–31.

21. Parker AJ, Smith JET, Krug K. Neural architectures for stereo vision. Philos Trans R Soc B Biol Sci. 2016;371(1697):20150261.

22. Yoshioka TW, Doi T, Abdolrahmani M, Fujita I. Specialized contributions of mid-tier stages of dorsal and ventral pathways to stereoscopic processing in macaque. Elife. 2021;10:1–76.

23. Backus BT, Fleet DJ, Parker AJ, Heeger DJ. Human cortical activity correlates with stereoscopic depth perception. J Neurophysiol. 2001;86(4):2054–68.

24. Preston TJ, Li S, Kourtzi Z, Welchman AE. Multivoxel pattern selectivity for perceptually relevant binocular disparities in the human brain. J Neurosci. 2008;28(44):11315–27.

25. Wang F, Yang W, Zhang L, Gundran A, Zhu X, Liu J, et al. Brain activation difference evoked by different binocular disparities of stereograms: An fMRI study. Phys Medica [Internet]. 2016;32(10):1308–13. Available from: 10.1016/j.ejmp.2016.07.007

26. Hutchison RM, Gallivan JP. Functional coupling between frontoparietal and occipitotemporal pathways during action and perception. Cortex. 2018;98:8–27.

27. Read JCA. Stereo vision and strabismus. Eye. 2015;29(2):214–24.

28. Singh P, Bergaal SK, Sharma P, Agarwal T, Saxena R, Phuljhele S. Effect of induced anisometropia on stereopsis and surgical tasks in a simulated environment. Indian J Ophthalmol. 2021;69(3):568.

29. Deepa BMS, Valarmathi A, Benita S. Assessment of stereo acuity levels using random dot stereo acuity chart in college students. J Fam Med Prim care. 2019;8(12):3850.

30. Chen X, Liao M, Jiang P, Sun H, Liu L, Gong Q. Abnormal effective connectivity in visual cortices underlies stereopsis defects in amblyopia. NeuroImage Clin [Internet]. 2022;34(November 2021):103005. Available from: 10.1016/j.nicl.2022.103005

31. Raichle ME, MacLeod AM, Snyder AZ, Powers WJ, Gusnard DA, Shulman GL. A default mode of brain function. Proc Natl Acad Sci. 2001;98(2):676–82.

32. Crossley NA, Mechelli A, Vértes PE, Winton-Brown TT, Patel AX, Ginestet CE, et al. Cognitive relevance of the community structure of the human brain functional coactivation network. Proc Natl Acad Sci [Internet]. 2013;110(28):11583–8. Available from: https://www.pnas.org/doi/abs/10.1073/pnas.1220826110

33. Friston K, Moran R, Seth AK. Analysing connectivity with Granger causality and dynamic causal modelling. Curr Opin Neurobiol [Internet]. 2013;23(2):172–8. Available from: 10.1016/j.conb.2012.11.010

34. Cohen JR, D’Esposito M. The segregation and integration of distinct brain networks and their relationship to cognition. J Neurosci. 2016;36(48):12083–94.

35. Horwitz B, Warner B, Fitzer J, Tagamets M-A, Husain FT, Long TW. Investigating the neural basis for functional and effective connectivity. Application to fMRI. Philos Trans R Soc B Biol Sci. 2005;360(1457):1093–108.

36. Thomas Yeo BT, Krienen FM, Sepulcre J, Sabuncu MR, Lashkari D, Hollinshead M, et al. The organization of the human cerebral cortex estimated by intrinsic functional connectivity. J Neurophysiol. 2011;106(3):1125–65.

37. Power JD, Schlaggar BL, Lessov-Schlaggar CN, Petersen SE. Evidence for hubs in human functional brain networks. Neuron [Internet]. 2013;79(4):798–813. Available from: 10.1016/j.neuron.2013.07.035

38. van den Heuvel MP, Sporns O. Rich-club organization of the human connectome. J Neurosci. 2011;31(44):15775–86.

39. Bertolero MA, Yeo BTT, D’Esposito M. The diverse club. Nat Commun [Internet]. 2017;8(1):1–10. Available from: 10.1038/s41467-017-01189-w

40. de Reus MA, van den Heuvel MP. Rich club organization and intermodule communication in the cat connectome. J Neurosci. 2013;33(32):12929–39.

41. Fornito A, Yoon J, Zalesky A, Bullmore ET, Carter CS. General and specific functional connectivity disturbances in first-episode schizophrenia during cognitive control performance. Biol Psychiatry. 2011;70(1):64–72.

42. Nicol RM, Chapman SC, Vértes PE, Nathan PJ, Smith ML, Shtyrov Y, et al. Fast reconfiguration of high-frequency brain networks in response to surprising changes in auditory input. J Neurophysiol. 2012;107(5):1421–30.

43. Kitzbichler MG, Henson RNA, Smith ML, Nathan PJ, Bullmore ET. Cognitive effort drives workspace configuration of human brain functional networks. J Neurosci. 2011;31(22):8259–70.

44. Lohia K, Soans RS, Raj D, Saxena R, Gandhi TK. Digital Stereo Test (DST): Static stereopsis assessment in simulated and real depth-deficit patients. J Vis [Internet]. 2023 Aug 1;23(9):4574. Available from: 10.1167/jov.23.9.4574

45. Ohlsson J, Villarreal G, Abrahamsson M, Cavazos H, Sjöström A, Sjöstrand J. Screening merits of the lang II, frisby, randot, titmus, and TNO stereo tests. J AAPOS. 2001;5(5):316–22.

46. Simons K. A comparison of the Frisby, Random-Dot E, TNO, and Randot circles stereotests in screening and office use. Arch Ophthalmol. 1981;99(3):446–52.

47. Neri P, Bridge H, Heeger DJ. Stereoscopic processing of absolute and relative disparity in human visual cortex. J Neurophysiol. 2004;92(3):1880–91.

48. Tsao DY, Vanduffel W, Sasaki Y, Fize D, Knutsen TA, Mandeville JB, et al. Stereopsis activates V3A and caudal intraparietal areas in macaques and humans. Neuron. 2003;39(3):555–68.

49. Carmon A, Bechtoldt HP. Dominance of the right cerebral hemisphere for stereopsis. Neuropsychologia. 1969;7(1):29–39.

50. Levi DM. Learning to see in depth. Vision Res [Internet]. 2022;200(March):108082. Available from: 10.1016/j.visres.2022.108082

51. Esteban O, Markiewicz CJ, Blair RW, Moodie CA, Isik AI, Erramuzpe A, et al. fMRIPrep: a robust preprocessing pipeline for functional MRI. Nat Methods [Internet]. 2019;16(1):111–6. Available from: 10.1038/s41592-018-0235-4

52. Gorgolewski K, Burns CD, Madison C, Clark D, Halchenko YO, Waskom ML, et al. Nipype: A flexible, lightweight and extensible neuroimaging data processing framework in Python. Front Neuroinform. 2011;5(August).

53. Dale AM, Fischl B, Sereno MI. Cortical Surface-Based Analysis. Neuroimage. 1999;9(2):179–94.

54. Friston KJ, Holmes AP, Price CJ, Büchel C, Worsley KJ. Multisubject fMRI studies and conjunction analyses. Neuroimage. 1999;10(4):385–96.

55. Glasser MF, Coalson TS, Robinson EC, Hacker CD, Harwell J, Yacoub E, et al. A multi-modal parcellation of human cerebral cortex. Nature [Internet]. 2016;536(7615):171–8. Available from: 10.1038/nature18933

56. Chen G, Glen DR, Saad ZS, Paul Hamilton J, Thomason ME, Gotlib IH, et al. Vector autoregression, structural equation modeling, and their synthesis in neuroimaging data analysis. Comput Biol Med [Internet]. 2011;41(12):1142–55. Available from: 10.1016/j.compbiomed.2011.09.004

57. Perry PO, Wolfe PJ. Null models for network data. 2012; Available from: http://arxiv.org/abs/1201.5871

58. Rubinov M, Sporns O. Complex network measures of brain connectivity: Uses and interpretations. Neuroimage [Internet]. 2010;52(3):1059–69. Available from: 10.1016/j.neuroimage.2009.10.003

59. Xue C, Sun H, Hu G, Qi W, Yue Y, Rao J, et al. Disrupted Patterns of Rich-Club and Diverse-Club Organizations in Subjective Cognitive Decline and Amnestic Mild Cognitive Impairment. Front Neurosci. 2020;14(October):1–14.

60. Breiman L, Friedman JH, Olshen RA, Stone CJ. Construction of trees from a learning sample. Classif Regres Trees, 1st ed; Taylor \& Fr Gr Abingdon, UK. 1984;21–3.

61. Horenstein C, Lowe MJ, Koenig KA, Phillips MD. Comparison of unilateral and bilateral complex finger tapping-related activation in premotor and primary motor cortex. Hum Brain Mapp. 2009;30(4):1397–412.

62. Koshino H, Minamoto T, Ikeda T, Osaka M, Otsuka Y, Osaka N. Anterior medial prefrontal cortex exhibits activation during task preparation but deactivation during task execution. PLoS One. 2011;6(8):e22909.

63. Brass M, von Cramon DY. The Role of the Frontal Cortex in Task Preparation. Cereb Cortex [Internet]. 2002;12(9):908–14. Available from: 10.1093/cercor/12.9.908

64. Minini L, Parker AJ, Bridge H. Neural modulation by binocular disparity greatest in human dorsal visual stream. J Neurophysiol. 2010;104(1):169–78.

65. Bridge H, Parker AJ. Topographical representation of binocular depth in the human visual cortex using fMRI. J Vis. 2007;7(14):1–14.

66. Cowey A, Porter J. Brain damage and global stereopsis. Proc R Soc London Ser B Biol Sci. 1979;204(1157):399–407.

67. Cowey A, Wilkinson F. The role of the corpus callosum and extra striate visual areas in stereoacuity in macaque monkeys. Neuropsychologia. 1991;29(6):465–79.

68. Bridge H. Effects of cortical damage on binocular depth perception. Philos Trans R Soc B Biol Sci. 2016;371(1697).

69. Janssen P, Vogels R, Liu Y, Orban GA. At least at the level of inferior temporal cortex, the stereo correspondence problem is solved. Neuron. 2003;37(4):693–701.

70. Thomas OM, Cumming BG, Parker AJ. A specialization for relative disparity in V2. Nat Neurosci. 2002;5(5):472–8.

71. Cottereau BR, McKee SP, Ales JM, Norcia AM. Disparity-tuned population responses from human visual cortex. J Neurosci. 2011;31(3):954–65.

72. Liu C, Li Y, Song S, Zhang J. Decoding disparity categories in 3-dimensional images from fMRI data using functional connectivity patterns. Cogn Neurodyn [Internet]. 2020;14(2):169–79. Available from: 10.1007/s11571-019-09557-6

73. Chen N, Chen Z, Fang F. Functional specialization in human dorsal pathway for stereoscopic depth processing. Exp Brain Res [Internet]. 2020;238(11):2581–8. Available from: 10.1007/s00221-020-05918-4

74. Iwami T, Nishida Y, Hayashi O, Kimura M, Sakai M, Kani K, et al. Common neural processing regions for dynamic and static stereopsis in human parieto-occipital cortices. Neurosci Lett. 2002;327(1):29–32.

75. Uka T, Tanabe S, Watanabe M, Fujita I. Neural correlates of fine depth discrimination in monkey inferior temporal cortex. J Neurosci. 2005;25(46):10796–802.

76. Janssen P, Vogels R, Orban GA. Three-dimensional shape coding in inferior temporal cortex. Neuron. 2000;27(2):385–97.

77. Verhoef BE, Bohon KS, Conway BR. Functional architecture for disparity in macaque inferior temporal cortex and its relationship to the architecture for faces, color, scenes, and visual field. J Neurosci. 2015;35(17):6952–68.

78. Georgieva S, Peeters R, Kolster H, Todd JT, Orban GA. The processing of three-dimensional shape from disparity in the human brain. J Neurosci. 2009;29(3):727–42.

79. Verhoef BE, Vogels R, Janssen P. Contribution of Inferior Temporal and Posterior Parietal Activity to Three-Dimensional Shape Perception. Curr Biol [Internet]. 2010;20(10):909–13. Available from: 10.1016/j.cub.2010.03.058

80. Janssen P, Vogels R, Liu Y, Orban GA. Macaque inferior temporal neurons are selective for three-dimensional boundaries and surfaces. J Neurosci. 2001;21(23):9419–29.

81. Roy JP, Komatsu H, Wurtz RH. Disparity sensitivity of neurons in monkey extrastriate area MST. J Neurosci. 1992;12(7):2478–92.

82. Maunsell JHR, Van Essen DC. The Connections of the Middel Temporal Visual Area (MT) and Their Relationship to a Cortical Hie. J Neurosci [Internet]. 1983;3(12):2563–86. Available from: http://www.jneurosci.org/content/jneuro/3/12/2563.full.pdf

83. DeAngelis GC, Newsome WT. Organization of disparity-selective neurons in macaque area MT. J Neurosci. 1999;19(4):1398–415.

84. Rutschmann RM, Greenlee MW. BOLD response in dorsal areas varies with relative disparity level. Neuroreport. 2004;15(4):615–9.

85. Uka T, DeAngelis GC. Linking neural representation to function in stereoscopic depth perception: Roles of the middle temporal area in coarse versus fine disparity discrimination. J Neurosci. 2006;26(25):6791–802.

86. Krug K, Parker AJ. Neurons in dorsal visual area V5/MT signal relative disparity. J Neurosci. 2011;31(49):17892–904.

87. Murphy AP, Leopold DA, Humphreys GW, Welchman AE. Lesions to right posterior parietal cortex impair visual depth perception from disparity but not motion cues. Philos Trans R Soc B Biol Sci. 2016;371(1697).

88. Lohia K, Soans RS, Saxena R, Gandhi TK. Disparity size changes functional integration during stereoscopic viewing: A pilot fMRI study. In: 29th Annual Meeting of The Organization for Human Brain Mapping (OHBM) [Internet]. Montréal, Canada; 2023 [cited 2023 Jul 27]. Available from: https://ww6.aievolution.com/hbm2301/index.cfm?do=abs.viewAbs&abs=1032

89. Kwee IL, Fujii Y, Matsuzawa H, Nakada T. Perceptual processing of stereopsis in humans: high-field (3.0-tesla) functional MRI study. Neurology. 1999;53(7):1599.

90. Paggetti G, Leff DR, Orihuela-Espina F, Mylonas G, Darzi A, Yang GZ, et al. The role of the posterior parietal cortex in stereopsis and hand-eye coordination during motor task behaviours. Cogn Process. 2015;16(2):177–90.

91. Poggio GF, Gonzalez F, Krause F. Stereoscopic mechanisms in monkey visual cortex: Binocular correlation and disparity selectivity. J Neurosci. 1988;8(12):4531–50.

92. Mendola JD, Lam J, Rosenstein M, Lewis LB, Shmuel A. Partial correlation analysis reveals abnormal retinotopically organized functional connectivity of visual areas in amblyopia. NeuroImage Clin. 2018;18:192–201.

93. Dai P, Zhang J, Wu J, Chen Z, Zou B, Wu Y, et al. Altered spontaneous brain activity of children with unilateral amblyopia: a resting state fMRI study. Neural Plast. 2019;2019.

94. Oishi H, Takemura H, Aoki SC, Fujita I, Amano K. Microstructural properties of the vertical occipital fasciculus explain the variability in human stereoacuity. Proc Natl Acad Sci. 2018;115(48):12289–94.

95. Forlim CG, Bittner L, Mostajeran F, Steinicke F, Gallinat J, Kühn S. Stereoscopic Rendering via Goggles Elicits Higher Functional Connectivity During Virtual Reality Gaming. Front Hum Neurosci. 2019;13(October).

96. Jolly JK, Sheldon AA, Alvarez I, Gallagher C, MacLaren RE, Bridge H. A low-cost telescope for enhanced stimulus visual field coverage in functional MRI. J Neurosci Methods [Internet]. 2021;350(September 2020):109023. Available from: 10.1016/j.jneumeth.2020.109023

