## Supplementary material for "Distinct rich and diverse clubs regulate coarse and fine binocular disparity processing: Evidence from stereoscopic task-based fMRI"

**Local graph features obtained using BCT across all regions and disparity conditions for right hemisphere (S1- S6)**

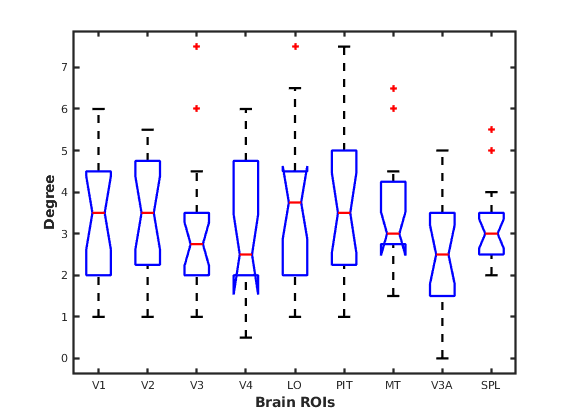

Fig. S1 Average degree of all regions of interest (ROIs) for 240 arc-sec condition. The red line in the middle of each box shows the median.

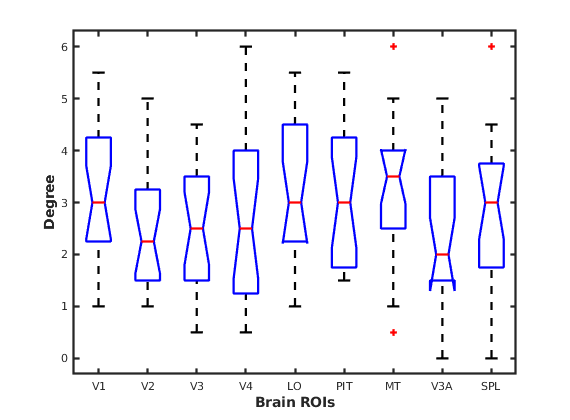

Fig. S2 Average degree of all ROIs for 480 arc-sec condition. The red line in the middle of each box shows the median.

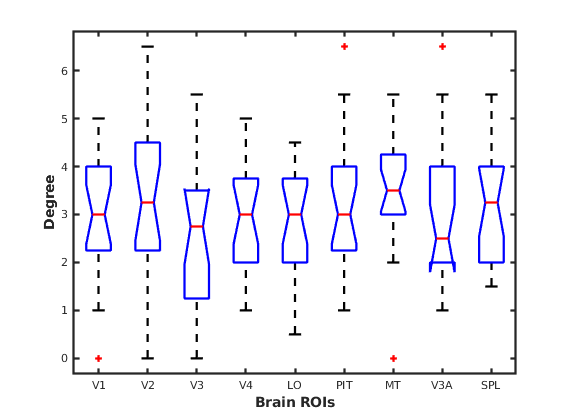

Fig. S3 Average degree of all ROIs for 800 arc-sec condition. The red line in the middle of each box shows the median.

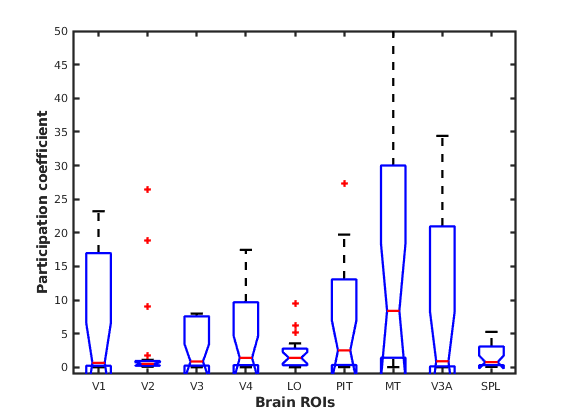

Fig. S4 Average participation coefficient of all ROIs for 240 arc-sec condition. The red line in the middle of each box shows the median.

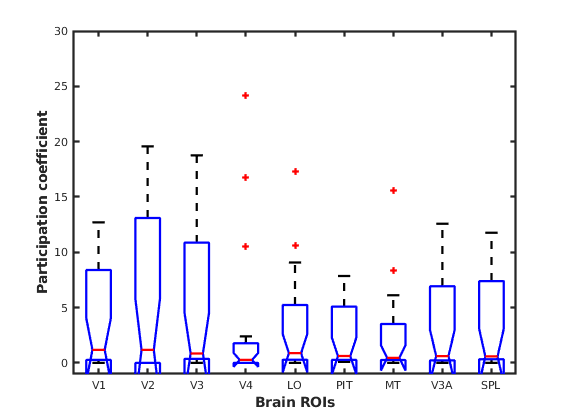

Fig. S5 Average participation coefficient of all ROIs for 480 arc-sec condition. The red line in the middle of each box shows the median.

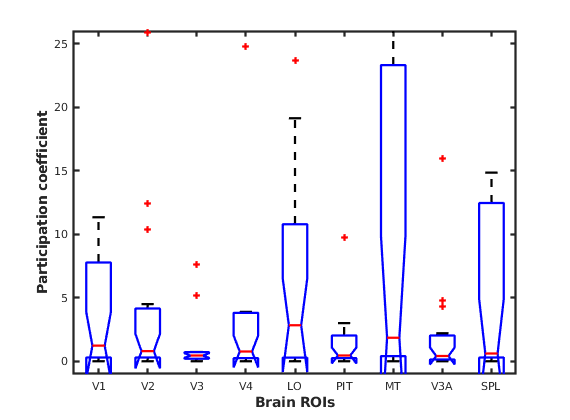

Fig. S6 Average participation coefficient of all ROIs for 800 arc-sec condition. The red line in the middle of each box shows the median.

**Local graph features obtained using BCT across all regions and disparity conditions for left hemisphere (S7- S12)**

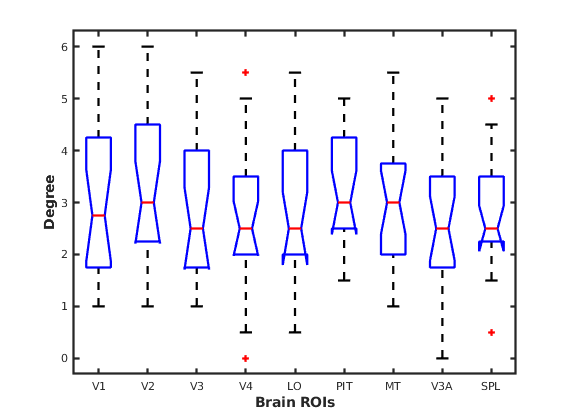

Fig. S7 Average degree of all regions of interest (ROIs) for 240 arc-sec condition. The red line in the middle of each box shows the median.

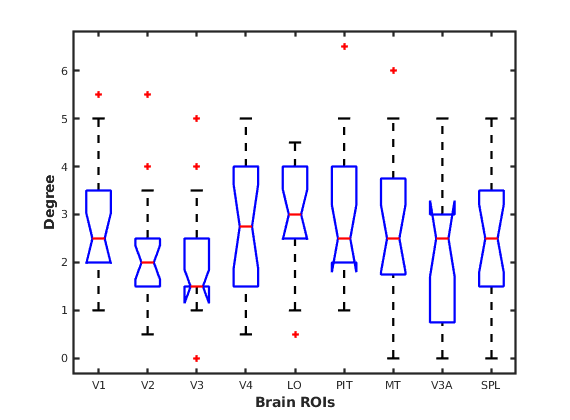

Fig. S8 Average degree of all ROIs for 480 arc-sec condition. The red line in the middle of each box shows the median.

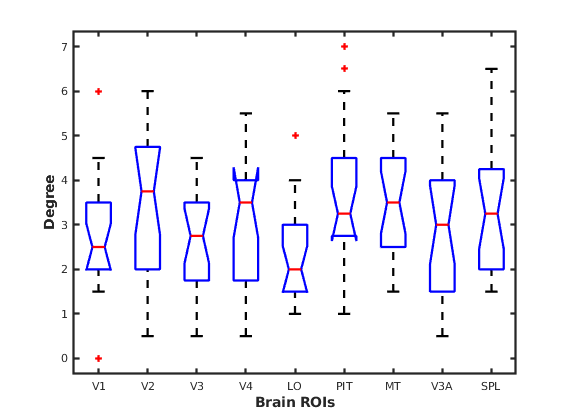

Fig. S9 Average degree of all ROIs for 800 arc-sec condition. The red line in the middle of each box shows the median.

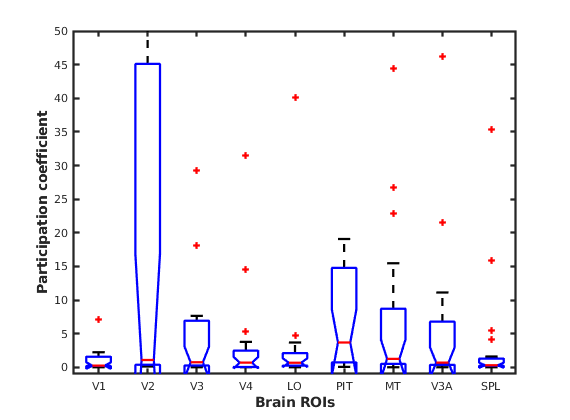

Fig. S10 Average participation coefficient of all ROIs for 240 arc-sec condition. The red line in the middle of each box shows the median.

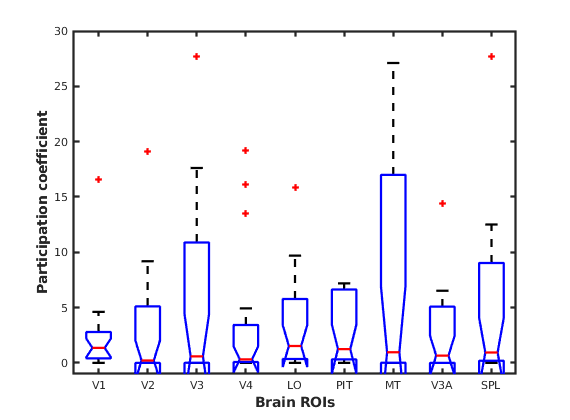

Fig. S11 Average participation coefficient of all ROIs for 480 arc-sec condition. The red line in the middle of each box shows the median.

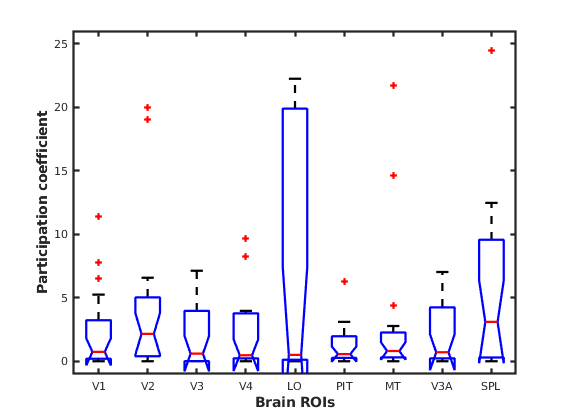

Fig. S12 Average participation coefficient of all ROIs for 800 arc-sec condition. The red line in the middle of each box shows the median.

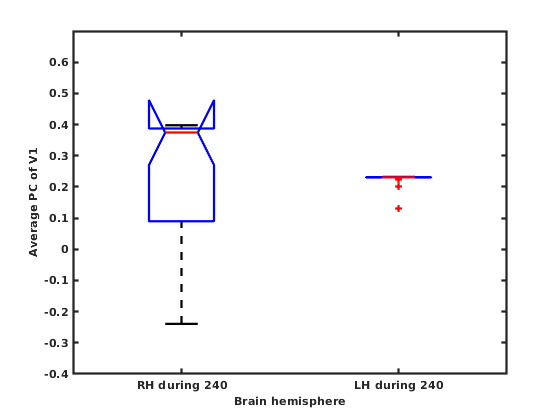

Fig. S13 Interhemispheric difference in Average Participation coefficient of V1 for 240 arc-sec condition. The red line in the middle of each box shows the median.

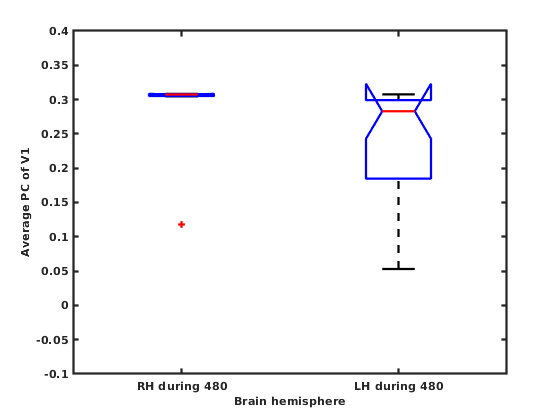

Fig. S14 Interhemispheric difference in Average Participation coefficient of V1 for 480 arc-sec condition. The red line in the middle of each box shows the median.

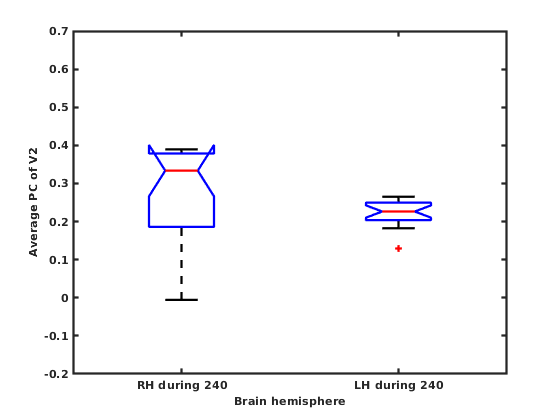

Fig. S15 Interhemispheric difference in Average Participation coefficient of V2 for 240 arc-sec condition. The red line in the middle of each box shows the median.

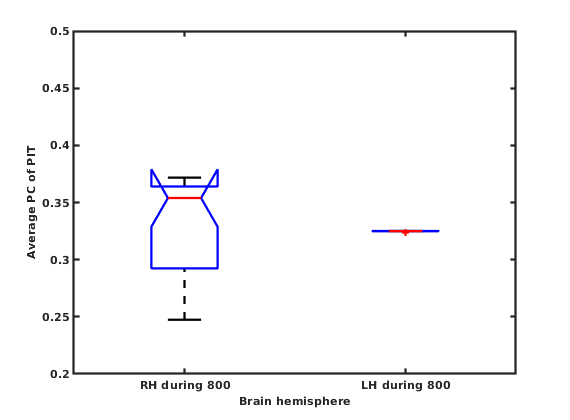

Fig. S16 Interhemispheric difference in Average Participation coefficient of PIT for 800 arc-sec condition. The red line in the middle of each box shows the median.

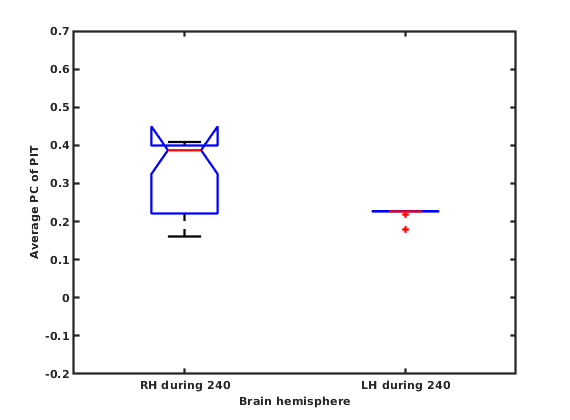

Fig. S17 Interhemispheric difference in Average Participation coefficient of PIT for 240 arc-sec condition. The red line in the middle of each box shows the median.

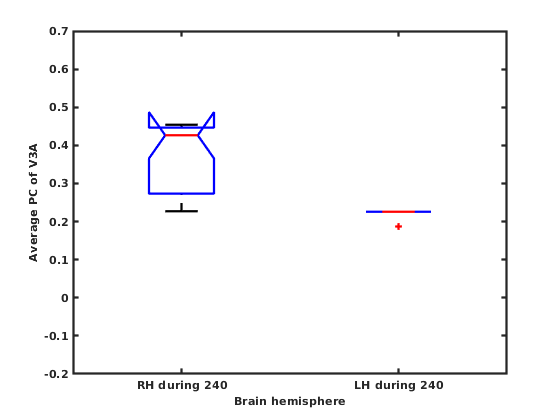

Fig. S18 Interhemispheric difference in Average Participation coefficient of V3A for 240 arc-sec condition. The red line in the middle of each box shows the median.

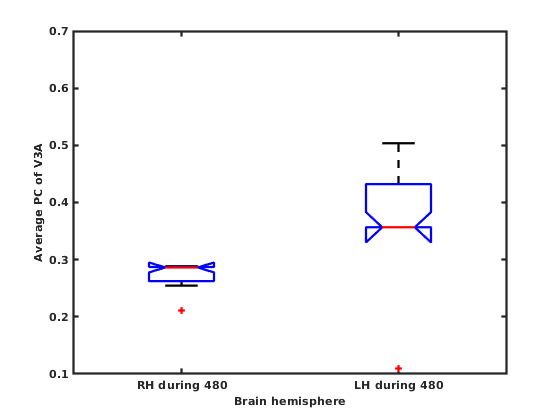

Fig. S19 Interhemispheric difference in Average Participation coefficient of V3A for 480 arc-sec condition. The red line in the middle of each box shows the median.

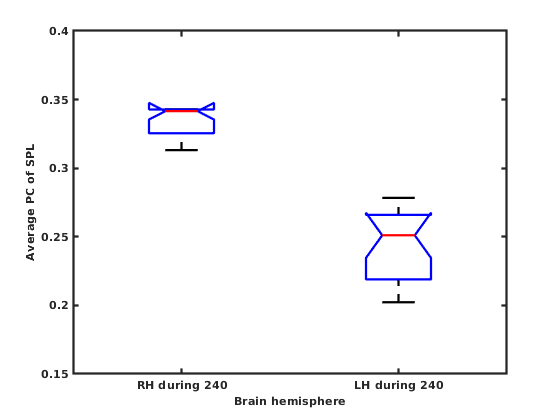

Fig. S20 Interhemispheric difference in Average Participation coefficient of SPL for 240 arc-sec condition. The red line in the middle of each box shows the median.

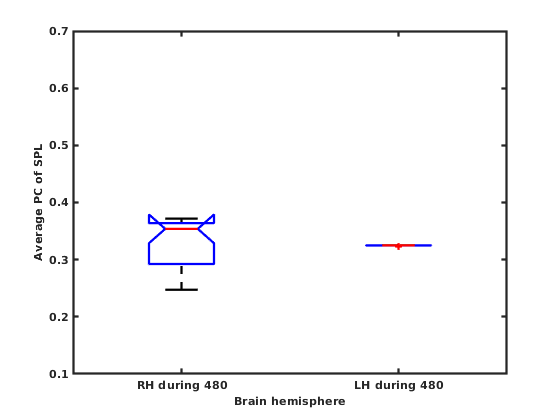

Fig. S21 Interhemispheric difference in Average Participation coefficient of SPL for 480 arc-sec condition. The red line in the middle of each box shows the median.

| **800 > SN** | **SNo.** | **#Voxels** | **CMx** | **CMy** | **CMz** | **Peak x** | **Peak y** | **Peak z** | **Alpha** |
| --- | --- | --- | --- | --- | --- | --- | --- | --- | --- |
|  | 1 | 193 | -19.1 | 96.9 | 12.6 | -11.5 | 100.5 | 13.5 | < 0.01 |
|  | 2 | 150 | -51.3 | 45.9 | 48.6 | -51.5 | 44.5 | 55.5 | < 0.01 |
|  | 3 | 125 | -18 | 81.7 | -11.5 | -23.5 | 80.5 | -16.5 | < 0.01 |
|  | 4 | 104 | 14.3 | 101.4 | 6.8 | 8.5 | 104.5 | 3.5 | < 0.01 |
|  | 5 | 99 | 53.5 | 53 | 51 | 52.5 | 50.5 | 57.5 | < 0.02 |
|  | 6 | 91 | 24.1 | 84 | -15.4 | 28.5 | 82.5 | -18.5 | < 0.02 |
|  | 7 | 42 | -59.5 | 48 | 31.2 | -61.5 | 48.5 | 31.5 | < 0.03 |
|  | 8 | 40 | -55.8 | 24.4 | -6.6 | -57.5 | 22.5 | -8.5 | < 0.03 |
| **480 > SN** | **SNo.** | **#Voxels** | **CMx** | **CMy** | **CMz** | **Peak x** | **Peak y** | **Peak z** | **Alpha** |
|  | 1 | 123 | 26.5 | 84.5 | -12.5 | 26.5 | 84.5 | -12.5 | < 0.02 |
|  | 2 | 81 | -11.5 | 94.5 | -8.5 | -11.5 | 94.5 | -8.5 | < 0.03 |
|  | 3 | 57 | -59.5 | 34.5 | 39.5 | -59.5 | 94.5 | 39.5 | < 0.03 |
|  | 4 | 47 | -23.5 | 76.5 | -10.5 | -23.5 | 76.5 | -10.5 | < 0.04 |
|  | 5 | 35 | 28.5 | 86.5 | -26.5 | 28.5 | 86.5 | -26.5 | < 0.05 |
|  | 6 | 34 | 20.5 | 100.5 | 7.5 | 20.5 | 100.5 | 7.5 | < 0.05 |
| **240 > SN** | **SNo.** | **#Voxels** | **CMx** | **CMy** | **CMz** | **Peak x** | **Peak y** | **Peak z** | **Alpha** |
|  | 1 | 156 | 10.5 | 90.5 | -16.5 | 10.5 | 90.5 | -16.5 | < 0.01 |
|  | 2 | 141 | 12.5 | 104.5 | 9.5 | 12.5 | 104.5 | 9.5 | < 0.01 |
|  | 3 | 105 | -27.5 | 96.5 | 13.5 | -27.5 | 96.5 | 13.5 | < 0.01 |
|  | 4 | 89 | -53.5 | 48.5 | 51.5 | -53.5 | 48.5 | 51.5 | < 0.01 |
|  | 5 | 64 | -37.5 | 70.5 | 53.5 | -37.5 | 70.5 | 53.5 | < 0.03 |
|  | 6 | 57 | 52.5 | 54.5 | 51.5 | 52.5 | 54.5 | 51.5 | < 0.03 |
|  | 7 | 46 | -11.5 | 90.5 | -10.5 | -11.5 | 90.5 | -10.5 | < 0.05 |
|  | 8 | 32 | 4.5 | -13.5 | 7.5 | 4.5 | -13.5 | 7.5 | < 0.04 |

Supplementary Table 1. Cluster table showing the clusters that survived the uncorrected cluster-forming p-threshold of p<0.001 (k = 39 for 800 > Noise, k=32 for 480 > Noise and k=29 for 240 > Noise) for overall alpha threshold of p<0.05 for all 3 conditions. Legend: #Voxels = No. of voxels in each cluster, k=cluster size threshold in voxels, alpha= Probability of a cluster > k, CM: Cluster of mass, Peak: coordinates of the peak voxel

| **SNo.** | **Region of Interest (ROI)** | **X (mm)** | **Y (mm)** | **Z (mm)** |
| --- | --- | --- | --- | --- |
| 1 | Left V1 | -7 | -89 | -6 |
| 2 | Left V2 | -12 | -94 | -15 |
| 3 | Left V3 | -14 | -93 | -19 |
| 4 | Left V4 | -23 | -91 | -21 |
| 5 | Left LO | -36 | -77 | 7 |
| 6 | Left PIT | -40 | -85 | -13 |
| 7 | Left MT | -48 | -68 | -13 |
| 8 | Left V3A | -21 | -91 | 23 |
| 9 | Left SPL | -34 | -47 | 57 |
| 10 | Right V1 | 8 | -86 | -5 |
| 11 | Right V2 | 19 | -91 | -13 |
| 12 | Right V3 | 16 | -74 | -12 |
| 13 | Right V4 | 52 | -69 | -3 |
| 14 | Right LO | 24 | -90 | 21 |
| 15 | Right PIT | 32 | -75 | 7 |
| 16 | Right MT | 45 | -63 | -13 |
| 17 | Right V3A | 10 | -90 | 31 |
| 18 | Right SPL | 30 | -46 | 53 |

Supplementary Table 2. Coordinates of Regions of interest (ROI) in MNI space. These ROIs are derived from the clusters in Supplementary Table 1 using Glasser HCP 2016 surface-based parcellation atlas across both hemispheres.

**Appendix: 1**

**Post-Experiment Questionnaire**

**Task: Static depth perception task**

Q1. Did you see a ⊔-like shape?    Yes	No

Q1a. If the answer to Q1 is yes, was the ⊔-like shape clearly visible to you?

Yes	No

Q2.  Did you see a ⊓-like shape? Yes	No

Q2a. If the answer to Q2 is yes, was the ⊓-like shape clearly visible to you?

Yes	No

Q3. Did you press the button each time you saw ⊓ or ⊓ shape?  Yes	No

Q3a. Can you think of an approximate number of times you pressed the button in the entire experiment? ……………

**(Note: 1 indicates least and 5 indicates most)**

Q4. On a scale of 1-5, how attentive were you in performing the experiment?

Task 1: 1	2	3	4	5

Q5. On a scale of 1-5, how much did you move while performing the experiment?

Task 1: 1	2	3	4	5

Q6.  Did you feel claustrophobic inside the scanner? Yes	No
